# Bi-channel Image Registration and Deep-learning Segmentation (BIRDS) for efficient, versatile 3D mapping of mouse brain

**DOI:** 10.1101/2020.06.30.181255

**Authors:** Xuechun Wang, Weilin Zeng, Xiaodan Yang, Chunyu Fang, Yunyun Han, Peng Fei

## Abstract

We have developed an open-source software called BIRDS (bi-channel image registration and deep-learning segmentation) for the mapping and analysis of 3D microscopy data of mouse brain. BIRDS features a graphical user interface that is used to submit jobs, monitor their progress, and display results. It implements a full pipeline including image pre-processing, bi-channel registration, automatic annotation, creation of 3D digital frame, high-resolution visualization, and expandable quantitative analysis (via link with Imaris). The new bi-channel registration algorithm is adaptive to various types of whole brain data from different microscopy platforms and shows obviously improved registration accuracy. Also, the attraction of combing registration with neural network lies in that the registration procedure can readily provide training data for network, while the network can efficiently segment incomplete/defective brain data that are otherwise difficult for registration. Our software is thus optimized to enable either minute-timescale registration-based segmentation of cross-modality whole-brain datasets, or real-time inference-based image segmentation for various brain region of interests. Jobs can be easily implemented on Fiji plugin that can be adapted for most computing environments.

## Introduction

The mapping of the brain and neural circuits is currently a major endeavor in neuroscience to understand fundamental and pathological brain processes[1–3]. Large projects which include the Mouse Brain Architecture project[4], the *Allen Mouse Brain Connectivity Atlas*[5] and the Mouse Connectome project map the mouse brain[6] in terms of cell types, long-range connectivity patterns and microcircuit connectivity. In addition to the large-scale collaborative, currently an increasing number of laboratories are also developing independent automated, or semi-automated framework for processing brain data obtained in specific projects[7–12]. With the improvement of experimental methods for dissection of connectivity and function, development of a standardized and automated computational pipeline to map, analyze, visualize and share brain data has become the major challenge to all such efforts[1, 7]. Thereby, the implementation of an efficient and reliable method is fundamentally required to define the accurate anatomical boundaries of brain structures, by which the anatomical positions of cells or neuronal connections could be determined to enable interpretation and comparison across experiments[10]. The commonly-used approach for automatic anatomical segmentation is to register the experimental image dataset with a standardized, fully segmented reference space, thus obtaining the anatomical segmentation for the experimental images[5, 8, 10, 13, 14]. There are currently several registration-based high-throughput image frameworks for analyzing large-scale brain datasets[7–10]. Most of these frameworks require the user to set a few parameters based on the image intensity or graphic outline, or completely convert the dataset into a framework-readable format to ensure the quality of the resulting segmentation. However, with the rapid development of sample labeling technology[15–17] and high-resolution whole-brain microscopic imaging[18–22], the obvious heterogeneous and non-uniform characteristics of brain structures make it difficult to use the traditional registration method to register datasets from different imaging platforms to a standard brain space with high accuracy. In this case, laborious visual inspection followed with manual correction are often required, which significantly reduce the productivity of the techniques. Therefore, we urgently need a robust, comprehensive registration method that can extract more unique features from the image data and provide accurate registration to different types of individual datasets.

Moreover, though registration-based methods can achieve full anatomical annotation in reference to a standard atlas for whole-brain datasets, their region-based 3D registration to whole-brain atlas lacks the flexibility to analyze incomplete brain datasets or a certain volume of interest[23], which is often the case in neuroscience research. Though some frameworks can register certain type of brain slab that contains complete coronal outlines in a way of slice by slice[7, 23–25], it remains very difficult to register small brain block without obvious anatomical outlines. As neural network has emerged as a technique of choice for image processing[26–29], deep learning-based brain mapping methods are also recently reported to directly provide segmentation/annotation of primary regions for 3D brain dataset[12, 30–33]. Such deep-learning-based segmentation networks are efficient in extracting pixel-level features, and thus are little dependent on the presence of global features, such as complete anatomical outlines, making it better suited for processing incomplete brain data, as compared to registration-based methods. On the other hand, the establishment of these networks still relies on sufficient amount of training dataset, which are laboriously registered, segmented and annotated. Therefore, a combination of image registration and neural network could possibly provide synergistic effect, and lead to more efficient and versatile brain mapping.

Here we provide an open source software as a Fiji[34] plugin, termed Bi-channel Image Registration and Deep-learning Segmentation (BIRDS), to support brain mapping efforts and feasible to analyze, visualize and share brain datasets. We have developed BIRDS to allow investigators to quantify and spatially map three-dimensional brain data in its own 3D digital space with reference to Allen CCFv3[35]. This facilitate the analysis in its native status at cellular level. The pipeline features 1. A bi-channel registration algorithm integrating the feature map with raw image data for co-registration with significantly improved accuracy achieved. 2. A mutually-beneficial strategy, in which the registration procedure can readily provide training data for network, while the network can efficiently segment incomplete brain data that are otherwise difficult for being registered with standardized atlas. The whole computational framework is designed to be robust and flexible, allowing its application to a wide variety of imaging systems (e.g., epifluorescence microscopy, light-sheet microscopy) and labeling approaches (e.g., fluorescent proteins, immunohistochemistry, and in situ hybridization). BIRDS pipeline offers a complete set of tool including image preprocessing, feature-based registration and annotation, visualization of digital map and quantitative analysis via link with Imaris, and neural network segmentation efficient for processing incomplete brain data. We demonstrate how BIRDS can be employed to fully automatic mapping of various brain structures and integrate multidimensional anatomical, neuronal labelling datasets. The whole pipeline has been packaged into a Fiji plugin, with a step-by-step tutorials provided to permit rapid implementation of the plugin in a standard laboratory environment.

## Results

### Bi-channel image registration with improved accuracy

Figure 1 shows our bi-channel registration procedure, which registers our experimental whole-brain images with standardized Allen institute mouse brain average template, and then provides segmentations and annotations from CCFv3 for experimental data. The raw high-resolution 3D image (1×1×10 μm^3^ voxel) obtained by serial two-photon tomography (STPT, see Methods) was first down-sampled into isotropic low-resolution data with 20-μm voxel size identical with the averaged Allen template image (Fig. 1a). It should be noted that, in addition to the individual difference, the sample preparation/mounting step could also lead to the deformation of samples, thereby posing extra challenge to the precise registration of experimental image to the averaged template (Fig. 1b, original data). To mitigate this issue before registration, we first subdivided the image stack into multiple sub-stacks (n=6 in our demonstration, Fig. 1b,) along the anterior-posterior (AP) axis (z axis) according to the landmarks identified in seven selected coronal planes referring to Allen Reference Atlas (Fig. 1a, Supplementary Fig. 1). Then we applied different re-slicing ratio to each sub-stack to finely rectify the deformation along AP axis. The down-sampled image stack obtained using this way shows higher similarity to the Allen template image, as compared to the raw experimental image stack (Fig. 1b). The implementation of such pre-processing step was beneficial to the better alignment of non-uniformly morphed brain data with the standardized template (Supplementary Fig. 4). We then applied a feature-based iterative registration of the Allen reference image with the pre-processed experimental images. It should be noted that previous registration methods are vulnerable to inadequate alignment accuracy[9, 10, 36], which is associated with inadequate registration information provided merely by the image data. To address this issue, here we used a phase congruency algorithm[37] to further extract the high-contrast edge and texture information from both the experimental and template brain images based on their relatively fixed anatomy features (Fig. 1c, Methods). Moreover, we also obtained the geometry features of both brains along their lateral-medial, dorsal-ventral, and anterior-posterior axes with enhanced axial mutual information (MI) extracted by a grayscale reversal processing[38, 39] (Fig. 1c, Supplementary Fig. 2, Methods). Thereby, an additional feature channel was formed based on these extracted feature information, and combined with the raw image channel to conduct our information-enriched bi-channel registration, which was proven to show notably better registration accuracy as compared to conventional single-channel registration methods (aMAP[9], ClearMap[10] and MIRACL[36]). During registration, through an iterative optimization of transformation from averaged Allen brain template to the experimental data, MI gradually reaches the maximum when the inverse grayscale images, PC images and the raw images are finally geometrically aligned (Fig. 1d). The displacement was presented in a grid form to illustrate the nonlinear deformation effect intuitively. The geometry wrapping parameters obtained from the registration process were then applied to the Allen annotation file to generate a transformed version specifically for the experimental data (Supplementary Fig.3). Our dual-channel registration has achieved fully automated registration/annotation at sufficiently high accuracy when processing STP experimental data of intact brain[40]. As for low-quality or highly deformed brain data (e.g., clarified brain with obvious shrinkage), though the registration accuracy of our method was accordingly reduced, it could still surpass previous methods obviously (Fig. 2). For such type of challenging data, we also developed an interactive graphic user interface (GUI) to readily permit manual correction of the visible inaccuracy in the annotation file, through finely tuning the selected corresponding points (Fig. 1e). Finally, an accurate 3D annotation could be generated and applied to the experimental data, either fully automatically (STP data) or after mild manual correction (LSFM data of clarified brain), as shown in Fig. 1f.

**Figure 1:**
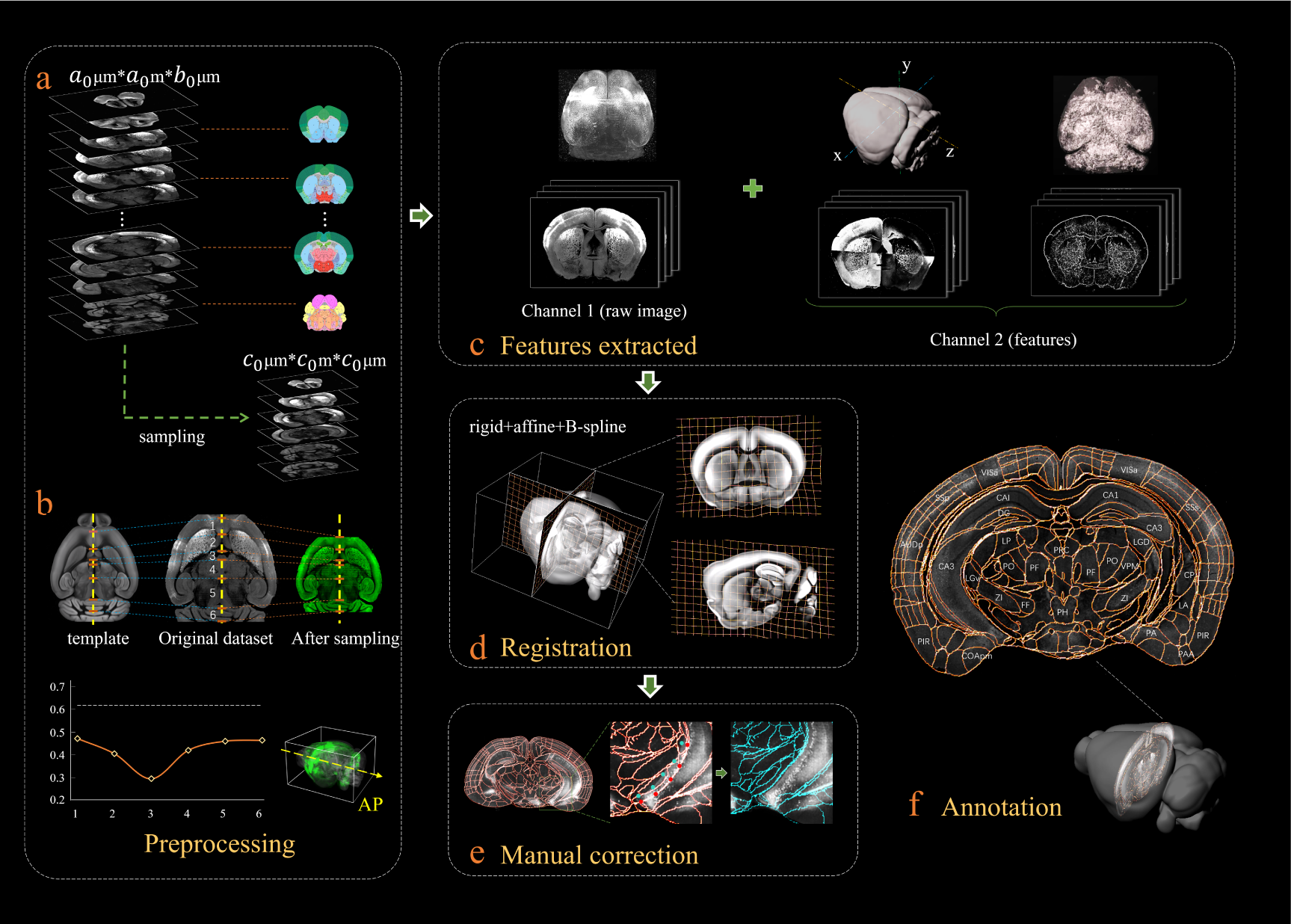
Bi-channel brain registration procedure. **(a)** Re-sampling of raw 3D image into an isotropic low-resolution one, which has the same voxel size (20 μm) with averaged Allen template image. **(b)** Dynamic down-sampling ratios applied to the raw experimental image along the AP axis (z-axis) stack by stack (indicated as 1 to 6). Each sub-stack was segmented according to the landmarks identified in seven selected coronal planes. This step can roughly restore the deformation of non-uniformly morphed sample data, thereby beneficial to the following registration with the Allen reference template. The plot shows the variation of the down-sampling ratio applied to the different z sub-stacks. **(c)** Additional feature channel containing a geometry and outline feature map extracted by grayscale reversal processing (left), as well as an edge and texture feature map extracted by phase congruency algorithm (right). This feature channel is combined with the raw image channel for implementing our information-enriched bi-channel registration, which shows improved accuracy as compared to conventional single-channel registration merely based on the raw images. **(d)** 3D view and anatomical sections (coronal and sagittal planes) of the registration results displayed in a grid deformed from the average Allen template. **(e)** Visual inspection and manual correction for the automatically-registered results of optically-clarified brain, which shows obvious deformation. With a GUI provided, this step can be readily operated by adjusting the interactive nodes on the annotation file (red points to light blue points). **(f)** The final atlas of experimental brain image containing region segmentations and annotations.

**Figure 2:**
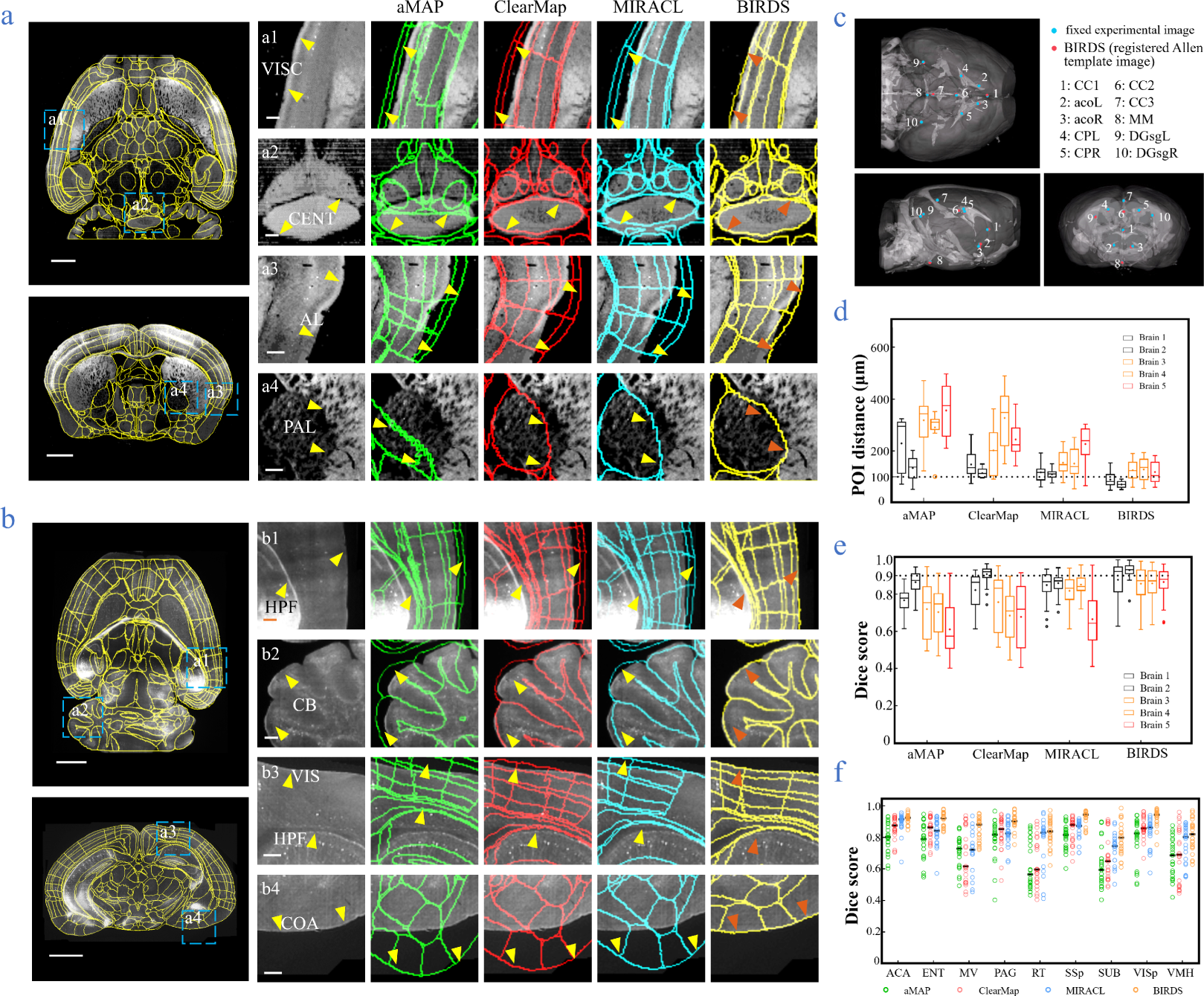
Comparison of BIRDS with conventional single-channel registration methods. **(a)** Comparative registration accuracy (STP data of intact brain) by four different registration methods, aMAP, ClearMap, MIRACL and our BIRDS. **(b)** Comparative registration accuracy (LSFM data of clarified brain) by four methods. Magnified views of four region-of-interests (**a1**-**a4**, **b1**-**b4**, blue boxes) selected from the horizontal (left, top) and coronal planes (left, bottom) are shown in the right four columns, to three-dimensionally detail the registration/annotation accuracy by each method. All comparative annotation results are directly outputted from the programs without manual correction. Scale bar, 1mm (whole-brain view), and 250 μm (magnified view). **(c)** Ten groups of 3D fiducial points of interest (POIs) manually identified across the 3D space of whole brains. The blue and red points belong to the fixed experimental image, and the registered Allen template image, respectively. The ten POIs are selected from following landmarks: *POIs*: *cc1*: corpus callosum, midline; *acoL*, *acoR*: anterior commisure, olfactory limb; *CPL, CPR*: Caudoputamen, Striatum dorsal region; *cc2*: corpus callosum, midline; *cc3*: corpus callosum, midline; *MM*: medial mammillary nucleus, midline; *DGsgL*, *DGsgR*: dentate gyrus, granule cell layer. The registration error by each method can be thereby quantified through measuring the Euclidean distance between each pair of POIs in the experimental image and template image. **(d)** Box diagram comparing the POIs distances of five brains registered by four methods. Brain 1, 2: STPT images of two intact brains. Brain 1 is also shown in **(a)**. Brain 3, 4, 5: LSFM images of three clarified brains (u-DISCO) which show significant deformations. Brain 5 is also shown in **(b)**. The median error distance of 50 pairs of POIs in the 5 brains registered by BIRDS is ~104 μm, which is compared to ~292 μm by aMAP, ~ 204 μm by ClearMap, and ~151 μm by most recent MIRACL. **(e), (f)** Comparative plot of Dice scores in nine registered regions of the five brains. The results are grouped by brains in **(e)**, and regions in **(f)**. The calculation is implemented at single nuclei level. When the results are analyzed by brains, BIRDS surpass the other 3 methods most on LSFM dataset #5, with 0.881 median Dice score being compared to 0.574 by aMAP, 0.72 by ClearMap, and 0.645 by MIRACL. At the same time, all the methods perform well on STP dataset #2 with median Dice of 0.874 by aMAP, 0.92 by ClearMap, 0.872 by MIRACL, and 0.933 by our BIRDS. When the results are compared by 9 functional regions, the median values acquired by our BIRDS were also higher than the other three methods. Even the lowest median Dice score by our method is still 0.799 (indicated by black line), which is notably higher than 0.566 by aMAP, 0.596 by ClearMap, and 0.722 by MIRACL.

### Comparison with conventional single-channel-based registration methods

Then we merged the experimental brain image with the registered annotation file to generate the 3D annotated image, and quantitatively compared its registration accuracy with aMAP, ClearMap, and MIRACL results. We made the comparison for both STPT data of intact brain that contained only minor deformation (Fig. 2a) and LSFM data of clarified brain (u-DISCO) that showed obvious shrinkage (Fig. 2b). It should be noted that here the annotated results of either previous single-channel methods or our bi-channel method are all from the automatic registration without any manual correction applied, and the averaged manual annotations by our experienced researchers serve as the ground truth for the quantitative comparisons. It’s visually obvious that, as compared to the other three methods (green: aMAP; red: ClearMap, blue: MIRACL in Fig. 2a, b), the Allen annotation files transformed and registered by our BIRDS method (yellow in Fig. 2a, b) are far better aligned with both STP (as shown in VISC, CENT, AL, and PAL regions, Fig. 2a) and LSFM (as shown in HPF, CB, VIS and COA regions, Fig. 2b) images. Furthermore, we manually labelled ten 3D fiducial points of interest (POIs) across the registered Allen template images together with their corresponding experimental images (Fig. 2c), and then measured the error distances between the paired anatomical landmarks in the two dataset so that the registration accuracy by each registration method could be quantitatively evaluated (Supplementary Fig. 5). As shown in Fig. 2d, the error distance distributions of POIs in five brains (two STP + three LSFM) registered by above-mentioned four methods were then quantified, showing the smallest median error distance (MED) was obtained by our method for all five brains. In two sets of STP data, only our BIRDS could provide MED below 100 μm (~80 μm, n=2), and this value slightly increased to ~120 μm for LSFM data (n=3), but still smaller than all the results by other three methods (aMAP, ~342 μm, n=3; ClearMap, ~258 μm, n=3; MIRACL, ~175 μm, n=3). Moreover, the Dice scores[41], defined as a similarity scale function used to calculate the similarity of two samples, for each method were also calculated at nucleus precision level based on nine functional regions in the five brains. The comparative results were grouped by brains and regions, as shown in Fig. 2e and f, respectively. The highest Dice scores with averaged median value > 0.89 (calculated by 5 brains, 0.75, 0.81, and 0.81 for aMAP, ClearMap, and MIRACL) or >0.88 (calculated by 9 regions, 0.74, 0.77, and 0.84 for aMAP, ClearMap, and MIRACL) were obtained by BIRDS, further confirming the best registration accuracy of our method. Through a comparative Wilcoxon test, our results are proven to be superior to other three methods (providing larger Dice scores) with P value<0.05 calculated either by brains, or by regions.

### Whole-brain digital map identifying the distributions of labeled neurons and axon projections

A 3D digital map (CCFv3) based on the abovementioned bi-channel registration was generated to support automatic annotation, analysis, and visualization of neurons in a whole mouse brain (Also see Methods). The framework thus enabled large-scale mapping of neuronal connectivity and activity, to reveal the architecture and function of brain circuits. Here, we demonstrated how BIRDS pipeline visualizes and quantifies the single-neuron projection patterns obtained by STPT imaging. A mouse brain containing six GFP labeled layer-2/3 neurons in the right visual cortex was imaged with STPT at 1×1×10-μm^3^ resolution[40]. After applying BIRDS procedure to the STPT image stack, we generated the 3D map of this brain (Fig. 3a). An interactive hierarchal tree of brain regions in the software interface could help navigate through correspond selected-and-highlighted brain region with its annotation information (Fig. 3b). Via linking with Imaris (Method), we visualized and traced each fluorescently labelled neuronal cell (n=5) across the 3D space of the entire brain (Fig. 3c).

**Figure 3:**
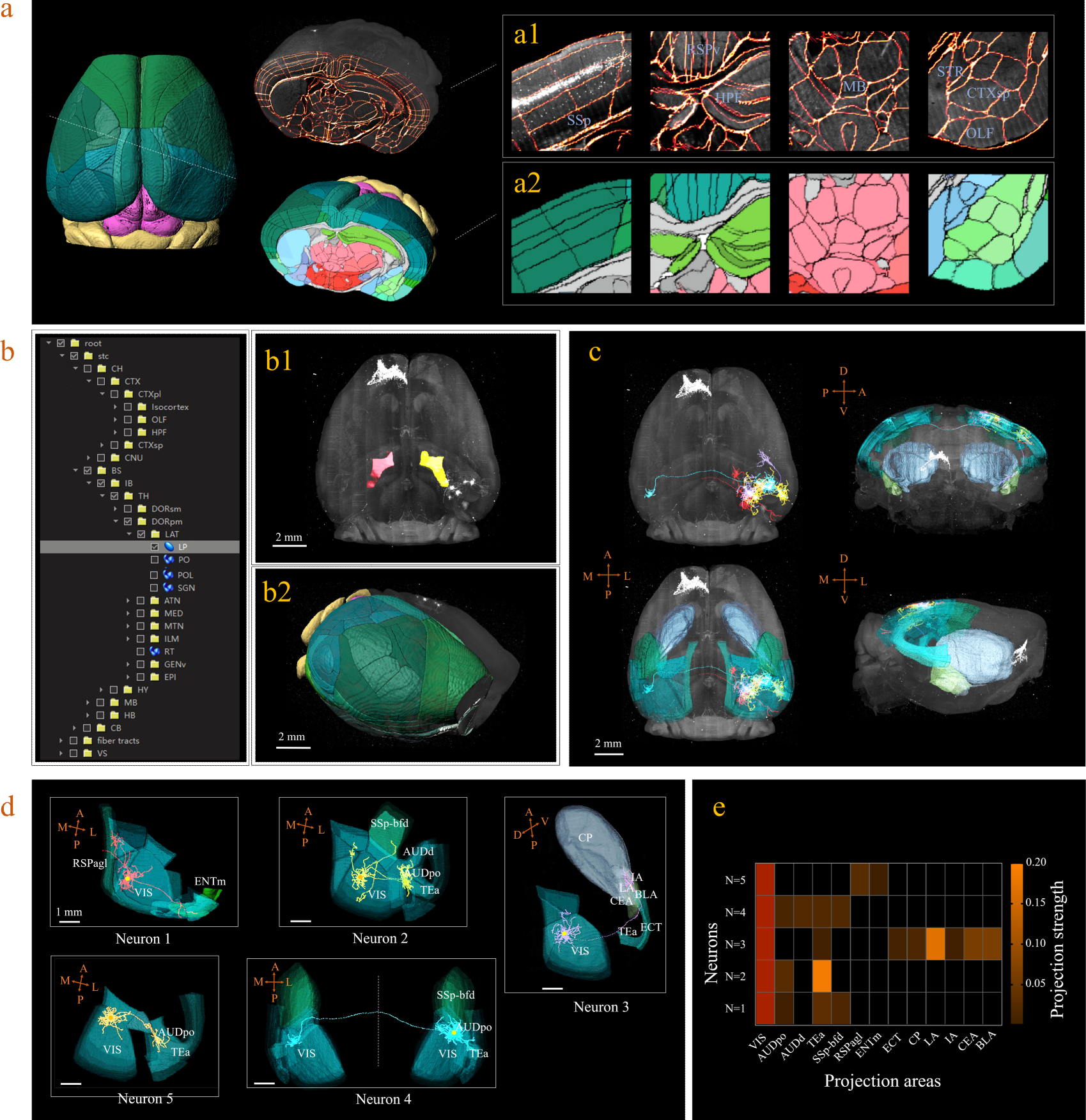
3D digital atlas of whole brain for visualization and quantitative analysis of inter-areal neuronal projections. **(a)** Rendered 3D digital atlas of the whole brain (a2, Pseudo color), which is generated from the registered template and annotation files (a1, overlay of annotation mask and image data). **(b)** Interactive hierarchical tree shown as a sidebar menu in the BIRDS program indexing the name of brain regions annotated in CCFv3. Clicking on any annotation name in the side bar of the hierarchal tree highlights the corresponding structure in 3D brain map (b1, b2), and vice versa. For example, brain region LP was highlighted in the space after its name was chosen in the menu (b1). 3D rendering of the individual brain after applying the deformation field in reverse to a whole brain surface mask. 3D digital atlas (CCFv3) is shown on the left; the individual brain is shown on the right (b2). **(c)** The distribution of axonal projections of five single neurons in 3D map space. The color-rendered space shown in horizontal, sagittal, and coronal views highlights multiple areas in the telencephalon, anterior cingulate cortex, striatum, and amygdala, which are all potential target areas of layer-2/3 neurons projections. **(d)** The traced axons of five selected neurons are shown in (N=5). ENTm, Entorhinal area, medial part, dorsal zone; RSPagl, Retrosplenial area, lateral agranular part; VIS, Visual areas; SSp-bfd, Primary somatosensory area, barrel field; AUDd, Dorsal auditory area; AUDpo, Posterior auditory area; TEa, Temporal association areas; CP, Caudoputamen; IA, Intercalated amygdalar nucleus; LA, Lateral amygdalar nucleus; BLA, Basolateral amygdalar nucleus; CEA, Central amygdalar nucleus; ECT, Ectorhinal area. **(e)** Quantification of the projection strength across the targeting areas of the five GFP-labelled neurons. The color code reflects the projections strengths of each neuron, determined as axon length per target area, normalized to the axon length in VIS.

BIRDS can be linked to Imaris to perform automated cell counting with higher efficiency and accuracy. Here we demonstrate it with an example brain where neurons were retrogradely labeled by CAV-mCherry injected to the right striatum and imaged by STPT at 1×1×10-μm^3^ resolution[40]. The whole-brain images stack was first processed by BIRDS to generate the 3D annotation map. Two of the example segregated brain areas (STR and BS) were outlined in the left panel of Fig. 4a. The annotation map and the raw image stack was then transferred to the Imaris which processed the images within each segregated area independently. Imaris calculated the local image statistics for cell recognition algorithm only using the image stack within each segregated area, therefore it fits the dynamic range of the local images to achieve the better result, shown in the middle column in the right panel of Fig. 4a. In contrast, the conventional Imaris automated cell counting program processed the whole-brain image stack at once to calculate the global cell recognition parameters for the every brain area, which can easily resulted in false positive or false negative counting in brain areas where the labeling signal was too strong or too weak compared to the global signal, as demonstrated in the STR and SB in the right column of the right panels of Fig. 4a respectively. BIRDS-Imaris program could perform automated cell counting for each brain area and reconstructed them over the entire brain. The 3D model of the brain-wise distribution of labeled striatum-projecting neurons was visualized by BIRDS-Imaris program as a 3D rendered brain image and projection views from three axes in Fig. 4b. BIRDS program could calculate the volume of each segregated region according to the 3D segregation map and the density of labeled cells across the brain as shown in Fig. 4c. Meanwhile, manual cell counting was also performed with every one out of four sections using ImageJ plugin (Fig. 4d). Compared to conventional Imaris result, the BIRDS-Imaris was more consistent to the manual one, especially for the brain regions where the fluorescent signal was at the high or low end of the dynamic range as highlighted with green boxes in Fig. 4e. Thanks to the 3D digital map generated by BIRDS pipeline, BIRDS-Imaris can process each segmented brain area separately, namely calculating the parameters for cell recognition algorithm using local image statistics instead of processing the whole-brain image stack at once. Such segmented cell counting strategy is much less demanding on computation resources, and moreover, it is optimized for each brain area to solve the problem that the same globe cell recognition parameter works poorly in certain brain regions with signal intensity at either of the two extreme ends of the dynamic range of the entire brain.

**Figure 4:**
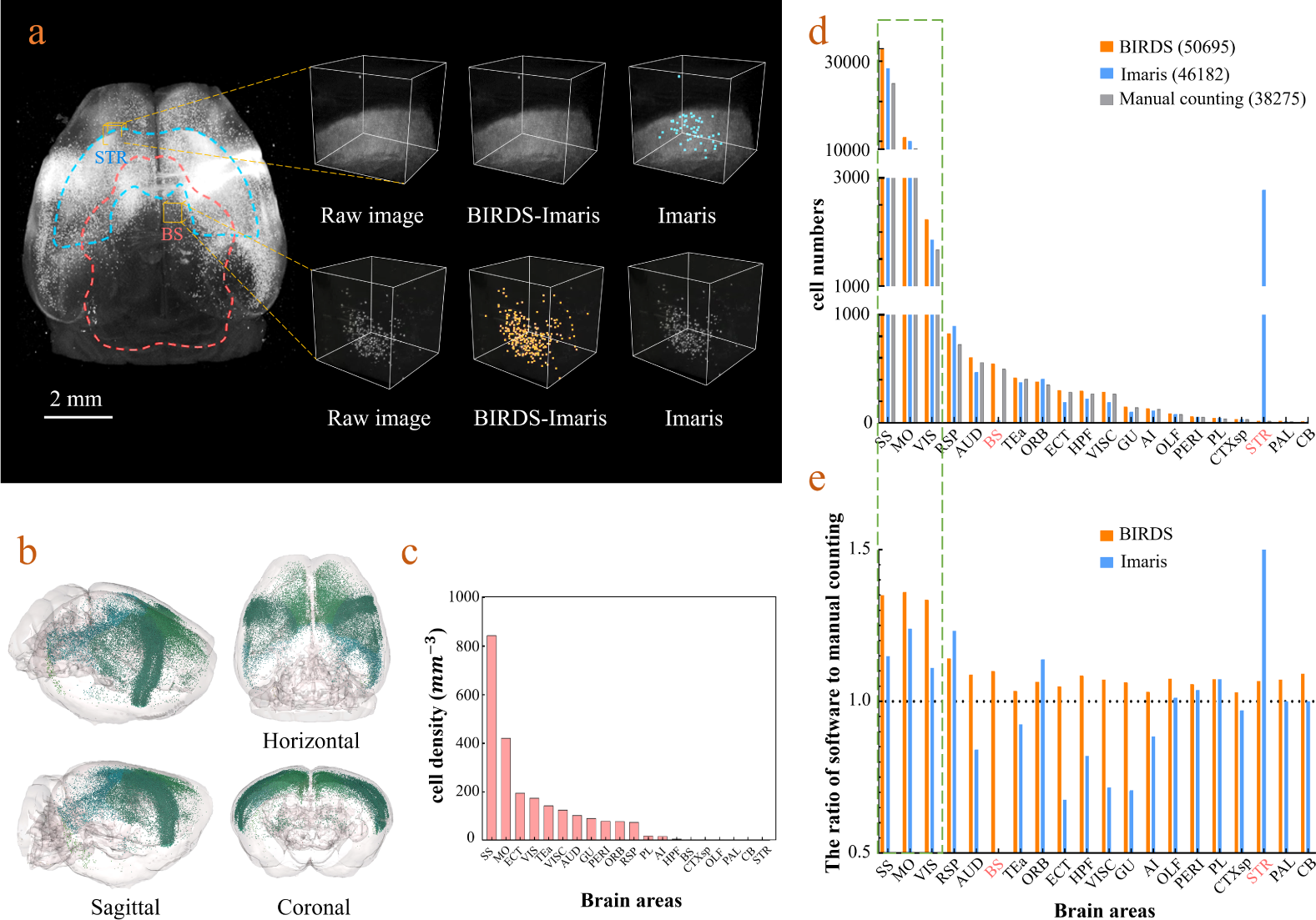
Cell-type-specific counting and comparison between different cell counting methods. **(a)** Cell counting of retrogradely labelled striatum-projecting cells. We selected two volumes (1×1×1 mm^3^) from SS and VIS areas, respectively, to show the difference in cell density and the quantitative results by BIRDS-Imaris and conventional Imaris. Here, separate quantification parameters set for different brain areas in BIRD-Imaris procedure lead to obviously more accurate counting results. Scale bar, 2 mm. **(b)** 3D-rendered images of labelled cells in the whole brain space, shown in horizontal, sagittal, and coronal views. The color rendering of the cell bodies is in accordance with the CCFv3, and the cells are mainly distributed in Isocortex (darker hue). **(c)** The cell density calculated for 20 brain areas. The cell densities of MO and SS are highest ((MO= 421.80 mm^−3^; SS=844.71 mm^−3^) among all the areas. GU, Gustatory areas; TEa, Temporal association areas; AI, Agranular insular area; PL, Prelimbic area; PERI, Perirhinal area; RSP, Retrosplenial area; ECT, Ectorhinal area; ORB, Orbital area; VISC, Visceral area; VIS, Visual areas; MO, Somatomotor areas; SS, Somatosensory areas; AUD, Auditory areas; HPF, Hippocampal formation; OLF, Olfactory areas; CTXsp, Cortical Subplate; STR, Striatum; PAL, Pallidum; BS, Brainstem; CB, Cerebellum. **(d)** Comparison of the cell numbers by three different counting methods, BIRDS, Imaris (3D whole brain directly), and manual counting (2D slice by slice for whole brain). **(e)** The cell counting accuracy by BIRDS-Imaris (orange) and conventional Imaris methods (blue), referring to the manual counting. Besides highly diverged accuracy for the 20 regions, the counting results by conventional Imaris in STR and BS regions are especially inaccurate.

### Inference-based segmentation of incomplete brain datasets using a deep learning procedure

In practice, the acquired brain dataset is often incomplete, owing to researcher’s particular interest to specific brain regions, or limited imaging conditions. The registration of such incomplete brain dataset to Allen template is often difficult due to the lack of sufficient morphology information for comparison in both datasets. To overcome this limitation, we further introduced a deep neural network (DNN) based method for efficient segmentation/annotation of incomplete brain sections with minimal human supervision. Herein, we optimized a Deeplab V3+ network, which is based on an encoding-decoding structure, for our deep-learning implementation (Fig. 5a). The input images passed through a series of feature processing stages in the network, with pixels being allocated, classified, and segmented into brain regions. It should be noted that the training of a neural network fundamentally requires a sufficiently large dataset containing various incomplete brain blocks which have been well segmented. Benefiting from our efficient BIRDS method, we could readily obtain a large number of such labelled data through cropping the processed whole brains and without experiencing time-consuming manual annotation. Various modes of incomplete brains, as shown in Fig. 5b, were hereby generated and sent to the DNN for iterative training, after which the skilled network could directly infer the segmentations/annotations for new modes of incomplete brain images (Methods). Here we validated the network performance on three different modes of input brain images cropped from the registered whole-brain dataset (STPT). The DNN has successfully inferred annotation results for cropped hemisphere, irregular cut of hemisphere, and a randomly cropped volume, as shown in Fig. 5c, d, e, respectively. The inferred annotations (red lines) have been found to be highly similar with the registered annotation results (green lines), in all three modes of incomplete data. To further quantify the inference accuracy, the Dice scores of the network-segmented regions were also calculated by comparing the network outputs with the ground truths, which are the registration result after visual inspection and correction (Supplementary Fig. 12). The averaged median Dice scores for the individual sub-regions in hemisphere, irregular cut of hemisphere, and random volume are 0.86, 0.87, and 0.87, respectively, showing a sufficiently high inference accuracy in most of brain regions, such as Isocortex, HPF, OLF, STR, etc. It’s noted that the performance of the network for segmenting the PAL, MBsta, P-sen regions remains sub optimal (Dice score 0.78 to 0.8), owing to their absences of obvious borders, and large structural variations across the planes (Supplementary Fig. 12). Finally, we applied the network inferences to generate the 3D atlases for these 3 incomplete brains, with segmenting the hemisphere into 18 regions as Isocortex, HPF, OLF, CTXsp, STR, PAL, CB, DORpm, DORsm, HY, MBsen, MBmot, MBsta, P-sen, P-mot, P-sat, MY-sen, MY-mot, the irregular cut of half telencephalon into 10 regions as Isocortex, HPF, OLF, CTXsp, STR, PAL, DORpm, DORsm, HY, MY-mot, and the random volume into 7 regions as Isocortex, HPF, STR, PAL, DORpm, DORsm, HY (Fig. 5f, g, h). Therefore, our DNN performs reasonably well even if the brain is highly incomplete. Furthermore, it can achieve the second-level fine segmentation within a small brain regions of interest. For example, we successfully segmented the hippocampus CA1, CA2, CA3, DG, as shown in Supplementary Fig. 13. Such a unique capability of network is possibly gained from the detection of pixel-level features rather than regions, and thereby substantially strengthens the robustness of our hybrid BIRDS method over conventional brain registration techniques when the data are highly incomplete/defective.

**Figure 5:**
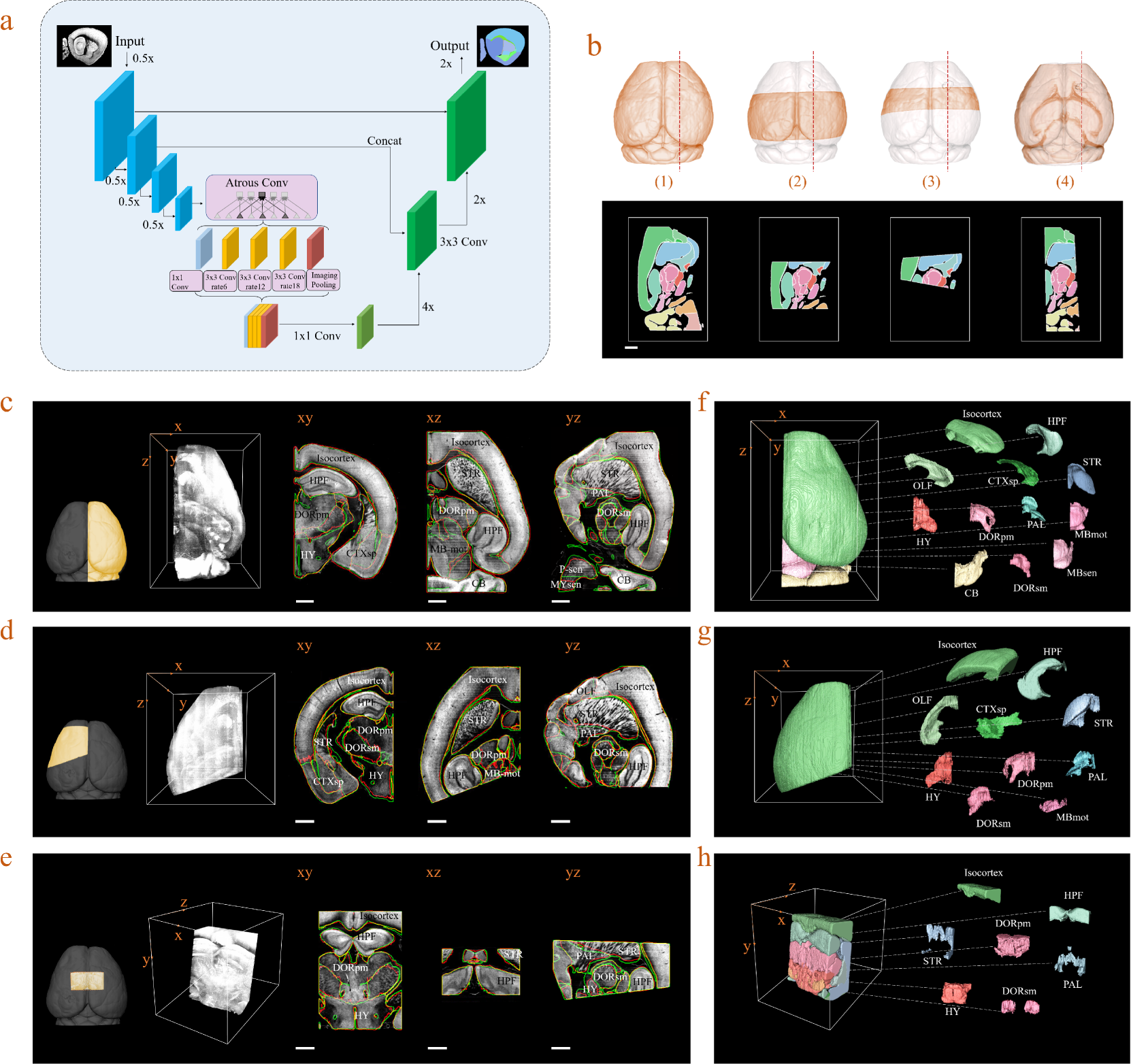
Inference-based network segmentation of incomplete brain data. **(a)** The deep neural network architecture for directly inferring the brain segmentation without registration required. The training datasets contain various types of incomplete brain images, which are cropped from the annotated whole-brain datasets created by our bi-channel registration beforehand. **(b)** Four models of incomplete brain datasets for network training: whole brain (1), a large portion of telencephalon (2), a small portion of telencephalon (3), and a horizontal slab of whole brain (4). Scale bar, 1 mm. **(c)-(e)** The inference-based segmentation results for three new modes of incomplete brain images as right hemisphere (**c**), an irregular cut of half telencephalon (**d**), and a randomly cropped volume (**e**). The annotated sub-regions are shown in x-y, x-z, and y-z planes, with Isocortex, HPF, OLF, CTXsp, STR, PAL, CB, DORpm, DORsm, HY, MBsen, MBmot, MBsta, P-sen, P-mot, P-sat, MY-sen, and MY-mot for right hemisphere, Isocortex, HPF, OLF, CTXsp, STR, PAL, DORpm, DORsm, HY, and MY-mot for irregular cut of half telencephalon, and Isocortex, HPF, STR, PAL, DORpm, DORsm, and HY for the random volume. Scale bar, 1mm. (**f**)-(**h**) The corresponding 3D atlases generated for these three incomplete brains.

## Discussion

In summary, we propose a bi-channel image registration method in conjunction with a deep-learning framework, to readily provide accuracy-improved anatomical segmentation for whole mouse brain in reference to Allen average template, and direct segmentation inference for incomplete brain dataset, which are otherwise not easy for being registered to standardized whole-brain space. The addition of brain feature channel to the registration process has greatly improved the accuracy of automatically registering individual whole-brain data with the standardized Allen average template. For processing various incomplete brain dataset, which are challenging to registration-based methods while remain very common in neuroscience research, we applied deep neural network to rapidly infer the segmentations. The sufficiently accurate results shown in different modes of incomplete data verify the advances of network segmentation. Though the full annotation by network is currently too computationally demanding as compared to registration-based segmentation, it’s undoubtedly a good complement to the registration-based segmentation. Therefore, in our hybrid BIRDS pipeline, the DNN inference greatly reinforced the inefficient side of registration while the registration also readily provided high-quality training data for DNN. We believe such a synergetic effect played in our method could provide a paradigm shift for enabling robust-and-efficient 3D image segmentation/annotation for biology research. With the unceasing development of deep learning, we envision the network-based segmentation will play more and more important role in the whole pipeline. A variety of applications, such as tracing of long-distance neuronal projections, and parallel counting of cell populations at different brain regions, are also enabled as a result of our efficient brain mapping. The BIRDS pipeline is now fully open-sourced by us, and also has been packaged into Fiji plugin to facilitate the potential biological researcher. We sincerely expect BIRDS method could immediately bring new insights for current brain mapping techniques, and thus allow us to further push the resolution and scale limits in the future exploration of brain space.

## Acknowledgements

We thank Yongsheng Zhang, Shaoqun Zeng, Haohong Li, Luoying Zhang, Man Jiang, Bo Xiong for discussions and comments on the work. Hao Zhang for the help on the code implementation. This work was supported by the National Key R&D program of China (2017YFA0700501 P.F.), the National Natural Science Foundation of China (21874052 for P.F.,31871089 for Y.H.), the Innovation Fund of WNLO (P.F.) and the Junior Thousand Talents Program of China (P.F. and Y.H.), the FRFCU (HUST:2172019kfyXKJC077 Y.H.).

## Author contributions

P.F. conceived the idea. P.F., and Y.H. oversaw the project. Y.H. and C.F. developed the optical setups and conducted the experiments. W.Z. and X.W. developed the programs. X.W., W.Z. and X.Y. processed, visualized and analyzed the data. X.W., W.Z., Y.H. and P.F. wrote the paper.

## Methods

### Acquisition of STPT image dataset

Brain 1 and 2 were obtained with STPT and each dataset encompassed ~90 Gigavoxels, e.g., 11980×7540×1075 in Dataset 1, with a voxel size of 1×1×10-μm^3^. The procedure of sample preparation and imaging acquisition were described in [39]. Briefly, the adult C57BL/6 mouse was anesthetized and craniotomy was performed on top of the right visual cortex. Individuals neuronal axons were labeled with pCAG-eGFP by two-photon microscopy guided single-cell electroporation and the brain was fixed by caridoperfusion of 4% PFA 8 days later. Striatum-projecting neurons were labeled by stereotactically injecting PRV-cre into the right striatum of tdTomato reporter mice (Ai14, JAX) and the brain was fixed cardioperfusion 30 days later. The brains were embedded in 5% oxidized agarose and imaged with a commercial STPT (TissueVision, USA) excited at 940 nm. Coronally, the brain was optically scanned every 10 μ m at 1 μm/pixel without averaging and physically sectioned every 50 μm. The power of excitation laser was adjusted to compensate the depth of optical sections.

### Acquisition of LSFM images dataset

Brain 3,4 and 5 were obtained with LSFM and each dataset encompassed ~400 Gigavoxels (~10000×8000×5000), with an isotropic voxel size of 1 μm^3^. 8-week-old Thy-GFP-M mice and the brain tissue was first clarified with uDISCO protocol[42] before imaging. Brain 3 and 4 were acquired using a custom-built Bessel plane illumination microscope, a type of LSFM modality employing non-diffraction thin Bessel light-sheet. Brain 5 was whole-brain 3D image of a Thy-GFP-M mice acquired using a lab-built selective plane illumination microscope (SPIM) [20], another LSFM modality combining Gaussian light-sheet with multi-view image acquisition/fusion. .

### Implementation of Bi-channel registration

#### Pre-processing of raw data

First, we developed an interactive GUI in the Fiji plugin to correspond the coronal planes in the Allen Reference Atlas (132 planes, 100-μm interval) with those in the experimental 3D image stack (e.g., 1075 planes with 10-μm stepsize in Dataset 1). As shown in Supplementary Fig. 1, seven coronal planes, from the anterior bulbus olfactorius to the posterior cerebellum, were identified across the entire brain, with their number of layers being recorded as *a*_*i*_ in template atlas, and *b*_*i*_ in the raw image stack. Therefore, *k* sub stacks (*k* = [1, 6], *k* ∊ *N*) was defined by these 7 planes (Fig. 1b). According to the ratio of step size between the Allen Reference Atlas (ARA) and its template image stack (100 μm to 20 μm), we also obtained the number of layers of the selected planes in the template image as *c*_*i*_ = 5*a*_*i*_ − 2. The reslicing ratio of the *k*^th^ sub-stack sandwiched by every two planes (a*k* to a*k+1*) was then calculated by: 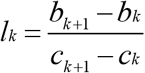. Each *l*_*k*_ was applied to the *k*^th^ sub-stack to obtain the resliced version of the sub-stack. Finally, the 6 resliced sub-stacks together formed a complete image stack of whole brain (20-μm stepsize), which had a rectified shape more similar to the Allen average template image, as compared to the raw experimental data. According to the isotropic voxel size of 20 μm in the template, the lateral size of voxel in the resliced image stack was also adjusted from originally 1 μm to 20 μm with a uniform lateral down-sampling ratio of 20 applied to all the coronal planes. The computational cost of abovementioned data resampling operation was low, taking merely ~5 minutes for processing 180 Gb raw STPT data on a Xeon workstation (E5-2630 V3 CPU).

#### Features extraction and processing

We extracted feature information based on the purified signals of the image data with backgrounds filtrated. We realized this through calculating the threshold values of the signals and backgrounds using by Huang’s fuzzy thresholding method[43], and removing the backgrounds according to the calculated thresholds. Then the feature information was detected using a phase congruency (PC) algorithm, which was robust to the intensity change of signals, and could efficiently extract corners, lines, textures information from the image. Furthermore, when the images had relatively low contrast at the border, which was very common in our study, the edge information could be much better retained using PC detection. Finally, the pixel intensity in the generated PC feature map can be calculated by following formula:

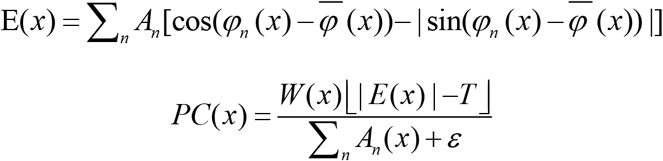

Where *x* is the angle vector of a pixel after Fourier transform of the image, E is the PC intensity, *T* is the noise threshold, *ε* is a small positive number (=0.01 in our practice) to prevent the denominator from leading to too large value of *PC*(*x*).

#### Multi-channel registration procedure

Image registration is fundamentally an iterative process optimized by a pre-designed cost function, which reasonably assesses the similarity between experimental and template datasets. Our bi-channel registration procedure is implemented based on the Elastix open-source program (Version 4.9.0)[44, 45]. Unlike conventional single-channel image registration, our method simultaneously registers all channels of multi-spectral input data using a single cost function defined as:

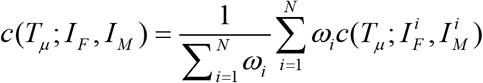

Where *N* represents the number of channels, and *ω*_i_ is the weight parameter of the channel. Since we used raw image stack, geometry feature map, and texture feature map for registration simultaneously, here *N*=3. 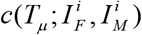 is the cost function of each channel, where 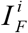 represents the fixed image (experimental data) and 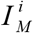 represents the moving image (template). *T*_*μ*_ denotes the deformation function of the registration model, with parameter *μ* being optimized during the iterative registration process. Here we used rigid+affine+B-spline three-level model for the registration, with rigid and affine transformations mainly for aligning the overall orientation differences between the datasets, and B-spline model mainly for aligning the local geometry differences. B-spline places a regular grid of control points onto the images. These control points are movable during the registration, and cause the surrounding image data to be transformed, thereby permitting the local, non-linear alignment of the image data. A stepsize of 30 pixels was set for the movement of control points in 3D space, and a five-levels coarse-to-fine pyramidal registration was applied for achieving faster convergence. During the iterative optimization of *T*_*μ*_, we used gradient descent method to efficiently approach the optimal registration of the template images to the fixed experimental images. The solution of *μ* can be expressed as

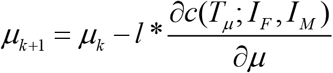

Where *l* is the learning rate, which also means the stepsize of the gradient descent optimization. The transformation parameters obtained from the multi-channel registration were finally applied to the annotation files to generate the atlas /annotation for the whole-brain data.

### Visualization and quantification of Brain-Map results

#### Generation of 3D digital map for whole brain

We obtained a low-resolution annotation (20-μm isotropic resolution) for entire mouse brain after registration. In order to generate a 3D digital framework based on the high-resolution raw image, *b*,*l* recorded in the down-sampling step were used for resolution restoration. Annotation information is to distinguish different brain regions by pixel intensity. In order to generate a digital frame for quantitative analysis, we introduced Marching cubes algorithm to generate 3D surface graphics, which is also, to generate the 3D digital maps. Then, through the programmable API link with Imaris, we could readily visualize the 3D digital map and perform various quantification in Imaris. After registration and interpolation applied to the experimental data, a 3D digital map was visualized in Imaris (9.0.0) invoked by our program. Then neuron tracing and cell counting tasks could be performed in Imaris at native resolution (e.g., 1×1×10 μm^3^ for STPT data). During the neural tracing process, the brain regions where the selected neurons passed through could be three-dimensionally displayed under arbitrary view. Furthermore, the cell counting could be performed in parallel by simultaneously setting a number of kernel diameters and intensity thresholds for different segmented brain regions.

#### Distance calculation

The Euclidean distance between one pair of landmarks shown in different image datasets (Fig. 2c) indicates the registration error and can be calculated as:

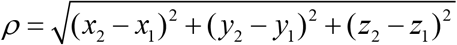

#### Calculation of Dice scores

Dice score is the indicator for quantifying the accuracy of segmentation, and can be calculated as:

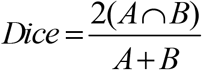

Where *A* is the ground-truth of segmentation while B is the result by Brain-Map. *A*∩*B* represents the number of pixels where *A* and *B* overlap, and *A*+*B* refers to the total number of pixels in *A* and *B*.

Here, with referring to BrainsMapi[8] methods, we compared the registration/segmentation results by four registration tools at both coarse region level and fine nuclei level. As we assessed the accuracy of these methods at brain-region level, ten brain regions, Outline, CB, CP, HB (hindbrain), HIP, HY (hypothalamus), Isocortex, MB (midbrain), OLF and TH (thalamus), were first selected from the entire brain for comparison. Then we further picked out 50 planes in each selected brain region (totally 500 planes for 10 regions), and manually segmented them to generate the reference results (ground truth). For nuclei-level comparison, we selected 9 small sub-regions, ACA, ENT, MV, PAG, RT, SSp, SUB, VISp, and VMH as targets, and performed similar operation on them with selecting five representative coronal sections for each region. To allow the manual segmentation as objective as possible, two skillful persons independently repeated the abovementioned process for 5 times, and a STAPLE algorithm[46] was used to fuse the 10 manual segmentation results to obtain the final averaged output as the ground-truth segmentation, for each region.

### Deep neural network segmentation

#### Generation of ground-truth training data

We chose 18 primary regions (level 4 and 5): Isocortex, HPF, OLF, CTXsp, STR, PAL, CB, DORpm, DORsm, HY, MBsen, MBmot, Mbsta, P-sen, P-mot, P-sat, MY-sen, and MY-mot, in whole brain for the DNN training and performance validation. The ground-truth annotation masks for these regions were readily obtained from our bi-channel registration procedure of BIRDS. For high-generosity DNN segmentation of whole brain and incomplete brain, we specifically prepared data training and validation as following:

Nine whole mouse brains containing 5000 annotated sagittal slices were first used as the training dataset. Then, the sagittal sections of whole brains were cropped, to generate different types of incomplete brains, as shown in Fig. 5b. The training dataset were thus comprised of both complete and incomplete sagittal sections (5000 and 4500 slices, respectively). The DNN trained by such datasets was able to infer segmentations for given complete or incomplete sagittal planes. Supplementary Fig. 9a compared the DNN segmentation results of both whole brain (a) and incomplete brains (b, c, d) with the corresponding ground-truths.

To demonstrate the performance of the DNN in segmenting sub-regions at finer scale, we chose the ground-truth images from coronal sections of specific hippocampus region in eight mouse brains (1100 slices in total). The corresponding ground-truth masks for the four major sub-regions of hippocampus, CA1, CA2, CA3, and the DG, were then generated by registration. We validated the DNN’ performance on segmenting these small sub-regions through comparison with the ground-truths at four different coronal planes.

#### Modified Deeplab V3+ network

Our DNN is based on the modification of Deeplab V3+, which contains a classical encoder-decoder structure[47] The main framework is based on Xception, which is optimized for deep separable convolutions and thereby reduces the computational complexity while maintains high performance. The network gradually reduces the spatial dimension of the feature map at the encoder stage, and allows complicated information to be easily outputted at deep level. At the final stage of the encoder structure, we introduce a dilated convolution Atrous Spatial Pyramid Pooling (ASPP), which increased the receptive field of convolution by changing the stepsize of atrous. The number of cores selected in ASPP is 6, 12, and 18. To solve the issue of aliased edges in inference results of conventional Deeplab V3+, we introduced more original image information into the decoder by serially convolving the 2× and 4× down-sampling results generated at the encoder stage and concatenating them to the decoder. The network was trained for ~30000 iterations with a learning rate of 0.001, learning momentum of 0.9 and output stride of 8. The training was implemented using two NVIDIA GeForce GTX 1080Ti graphics cards, and took approximately 40 hours.

