## supplementary file for "Bi-channel Image Registration and Deep-learning Segmentation (BIRDS) for efficient, versatile 3D mapping of mouse brain"

### Supplementary Information

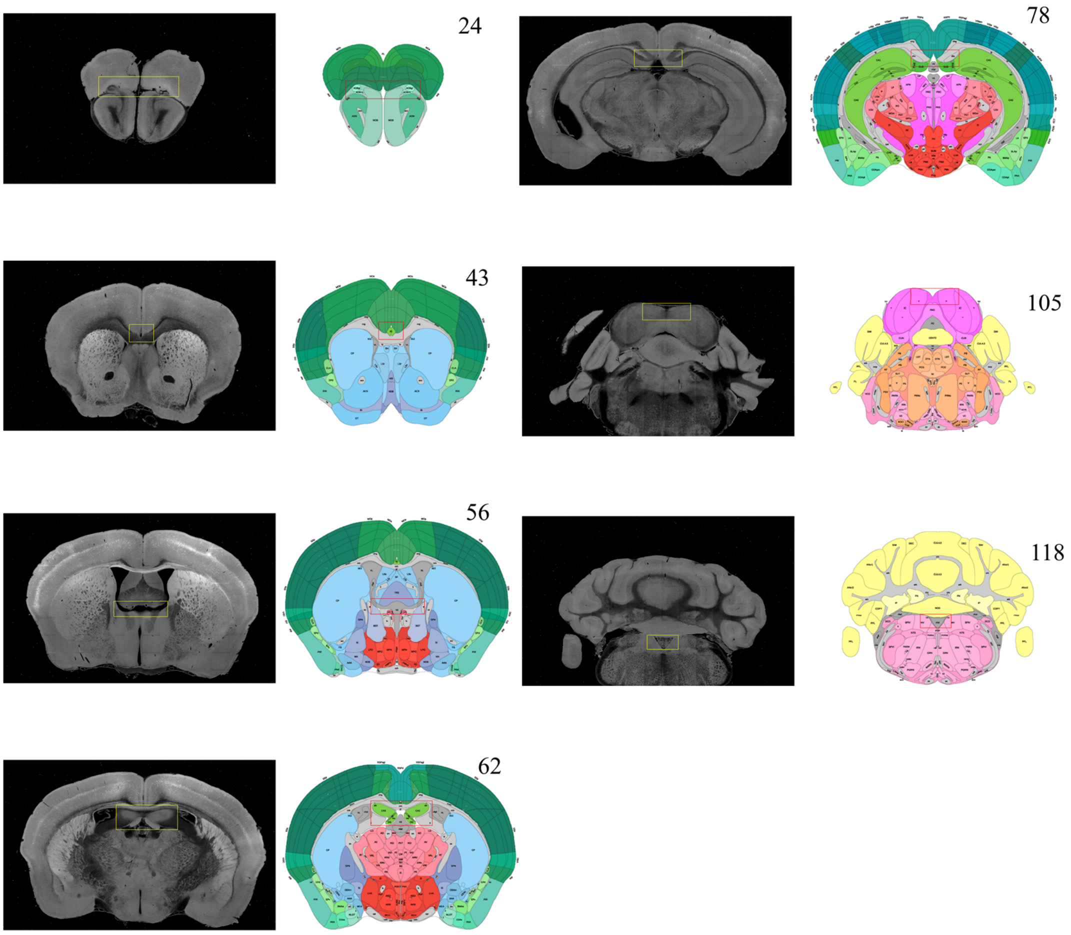


**Supplementary Figure 1: Selection of featured coronal planes corresponded to the Allen Reference Atlas, for subdividing the entire brain volume into multiple sub-stacks along AP axis.** The GUI of BIRDS program first permits the selection of a series of coronal planes (monochrome images) from the deformed 3D brain data, which are corresponded to the Allen Reference Atlas (#23, 43, 56, 62, 78, 105, and 118 of total 132 slices). The selection of coronal planes corresponded to the Allen Reference Atlas is based on the identification of several anatomical features, as shown in the paired yellow-red rectangular boxes. Then the 6 sub-stacks segmented according to these 7 planes are processed with different down-sampling ratios (from 0.297 to 0.48, Fig. 1b), to obtain the corresponding rectified stacks with 1:1 scale ratio to the Allen template images (total 660 slices). These rectified sub-stacks are finally combined to form an entire 3D brain volume with restored shape more similar to the Allen average template.

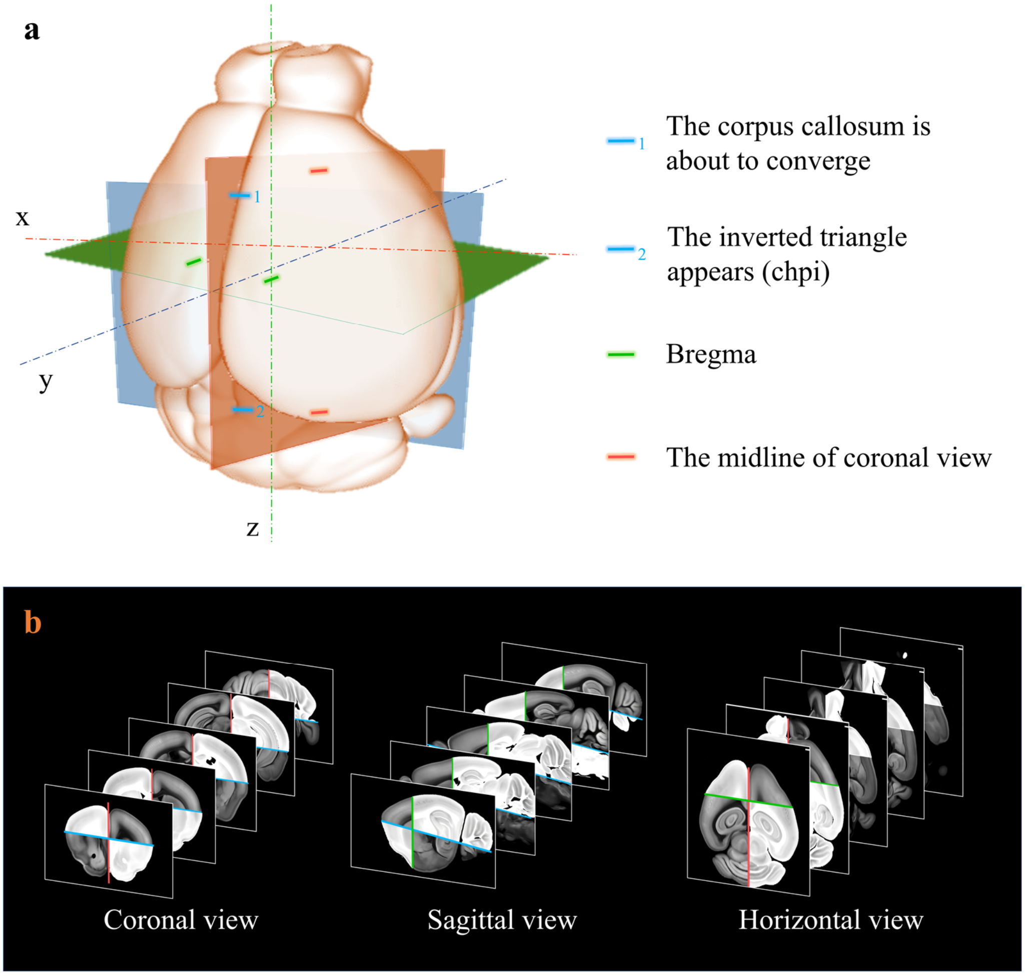


**Supplementary Figure 2: Extraction of axes features for registration. a** Determination of sagittal (orthogonal to lateral-medial axis, orange); horizontal (orthogonal to dorsal-ventral axis, blue); and coronal (orthogonal to anterior-posterior axis, green) planes based on the identification of specific anotomical features, as shown in a. **b** Grayscale reversal processing applied to the background-filtrated image dataset, according to the above-defined axis planes. The extraction of pure signals together with artificial reversal operation generate an extra feature map containing the geometry information of the axes, which is benefical to the improvement of registration accuracy.

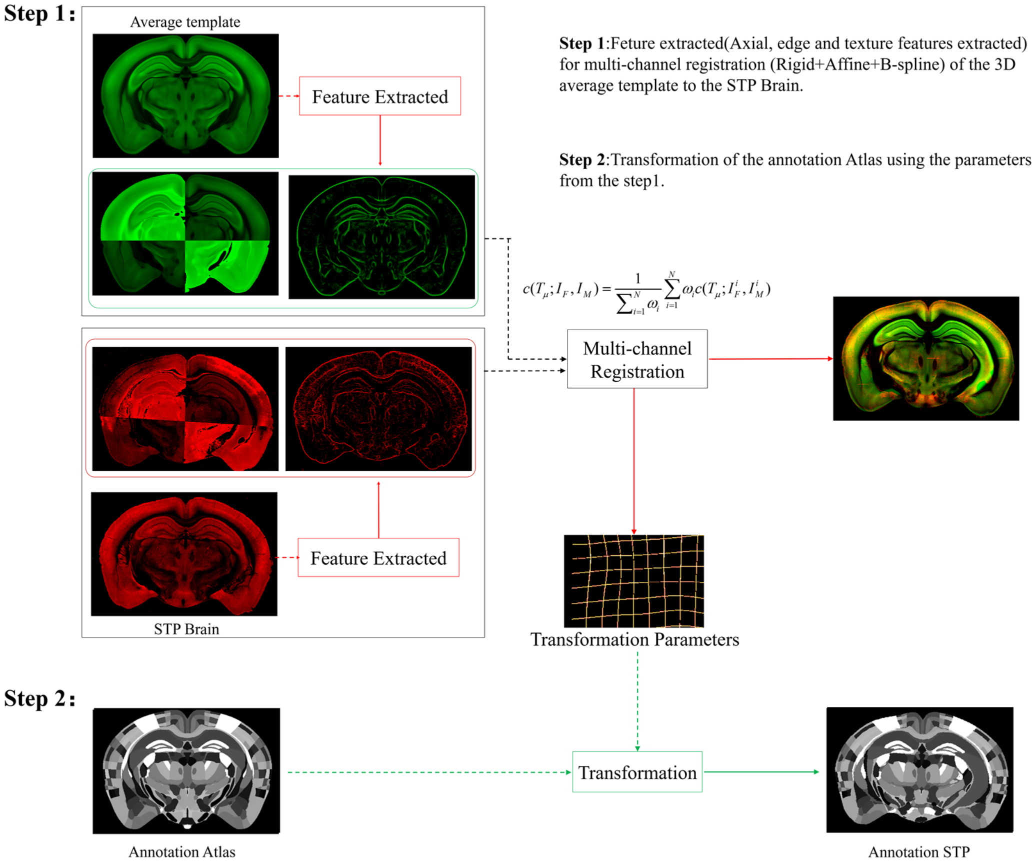


**Supplementary Figure 3: Schematic of the multi-channel registration and annotation process.** For both experimental and template data, an additional feature channel containing texture features extracted by PC and axial geometry features added by grayscale reversal, are combined with the raw image channel for our multi-channel registration based on rigid, affine and B-spline transformation. This procedure simultaneously registers all the multi-spectral input data using a single cost function, as shown in Step 1. The transformation parameters obtained from the image registration procedure are then applied to the template mouse annotation atlas (Allen Institute, CCF v3) to transform the template annotation file into an individual one that specifically fits our experimental brain image, and thus describes the anatomical structures in the whole brain space (Step 2).

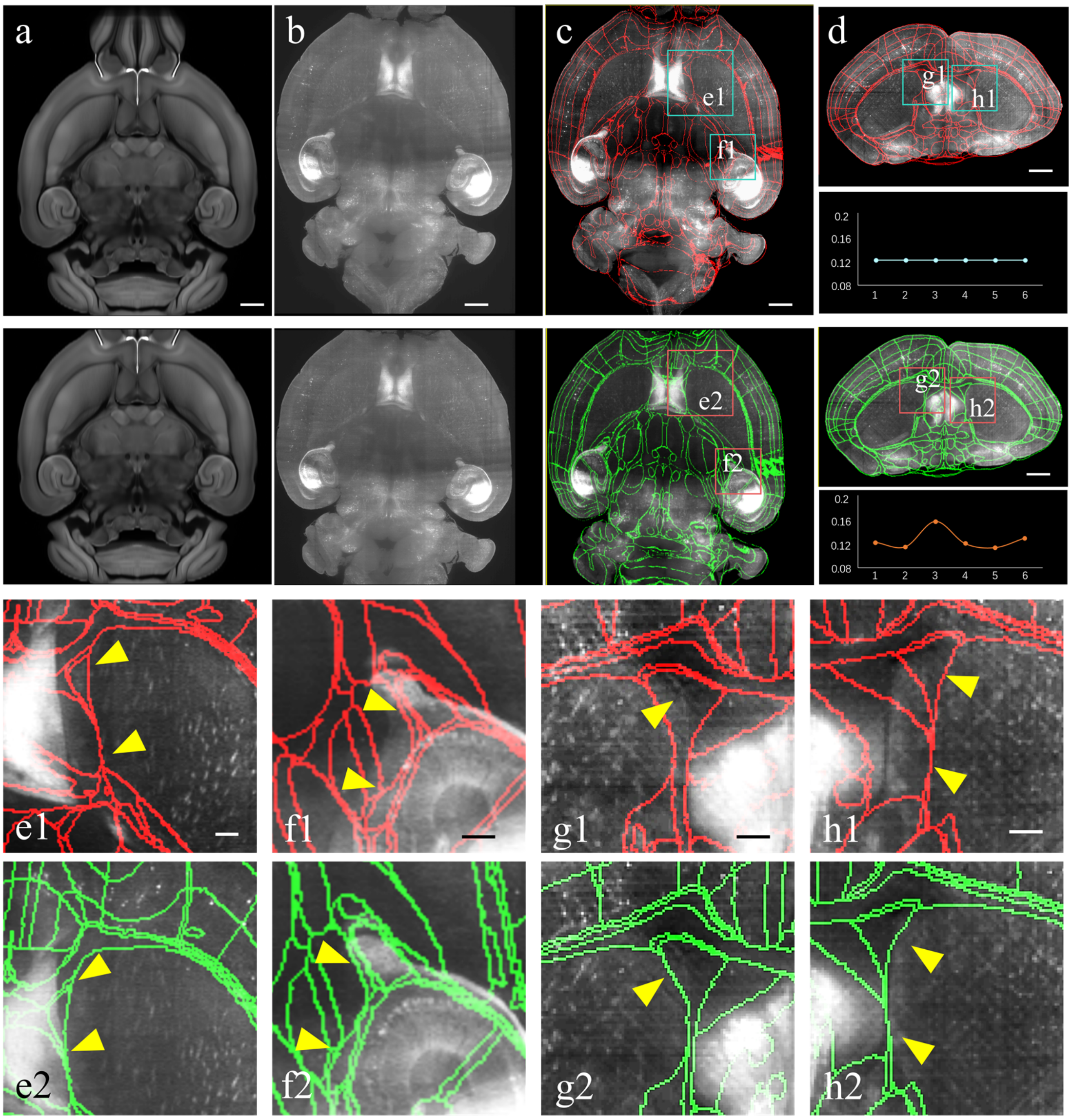


**Supplementary Figure 4: Improved registration accuracy enabled by the above mentioned shape rectification preprocessing. a-d** Automatical registration results of the experimental brain data applied with conventional uniform and our non-uniform downsampling, respectively. From the **a** to **d**, we comparatively show the the moving template image, fixed experimental image, corresponding segmentation results in horizontal and coronal views, respectively. Scale bars, 1mm. **e, f** Magnified views of 2 region-of-interests (boxes) shown in the registered-and-annotated horizontal planes **c**. The yellow arrows indicate the improved accuracy by our BIRDS method **e2**, **f2**, as compared to conventional uniform downsampling registration **e1**, **f1**. **g, h** Magnified views of 2 region-of-interests (boxes) shown in the registered-and-annotated coronal planes **d**. The yellow arrows again indicate the improved accuracy by our method **g2**, **h2**, as compared to conventional uniform downsampling registration **g1**, **h1**. Scale bar, 250 μm.

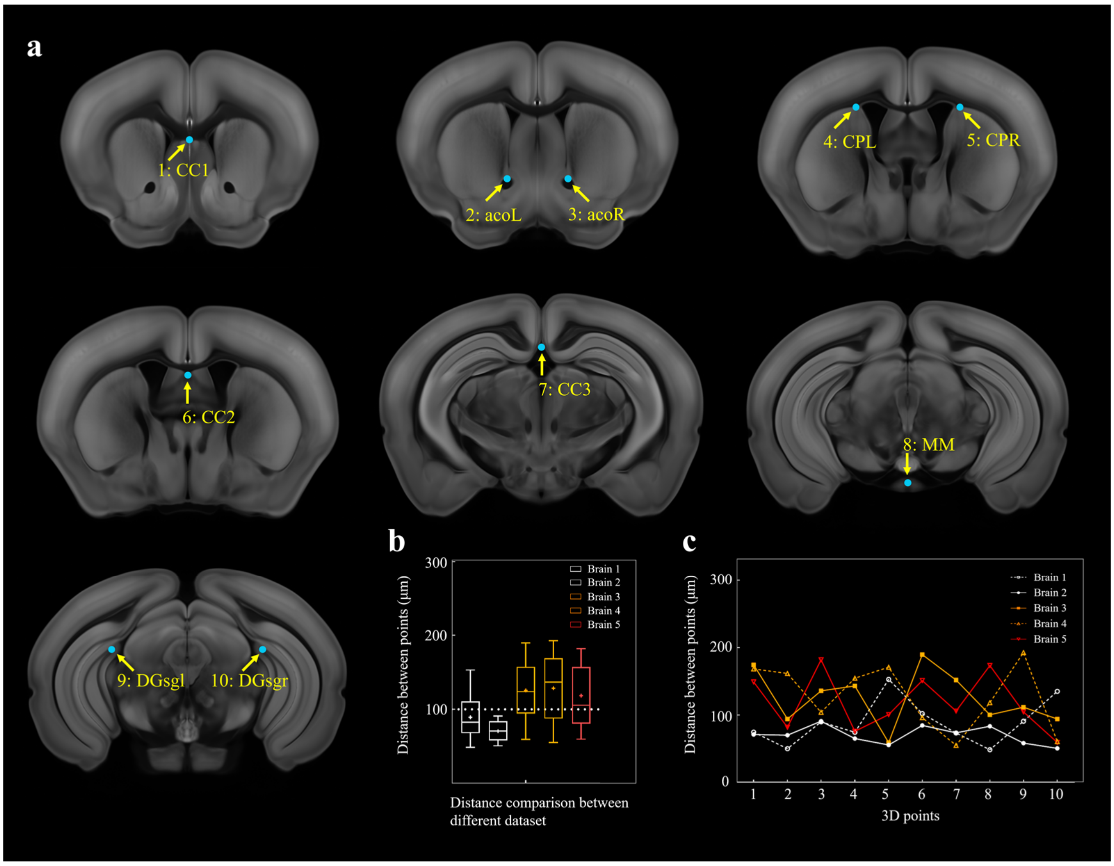


**Supplementary Figure 5: Selection of ten fiducial points of interest (POIS) for measuring the error distance between registered brains. a** Ten selected feature points in Allen template image. (1) cc2: corpus callosum, midline. (2),(3) acoL, acoR: anterior commisure, olfactory limb. (4),(5) CPL , CPR: Caudoputamen, Striatum dorsal region. (6) cc3: corpus callosum, midline. (7) cc1: corpus callosum, midline. (8) MM: medial mammillary nucleus, midline. (9),(10) DGsgL, DGsgR: dentate gyrus, granule cell layer. **b** Box diagrams showing the error distances between the paired POIs in registered template and experimental brains (n=5 brains, which are also used for **Fig. 2**). Using BIRDS method, the median error distance of total 50 pairs of POIs in 5 brains is ~104 μm. **c** Line plot further showing the distance between each pair of POI.

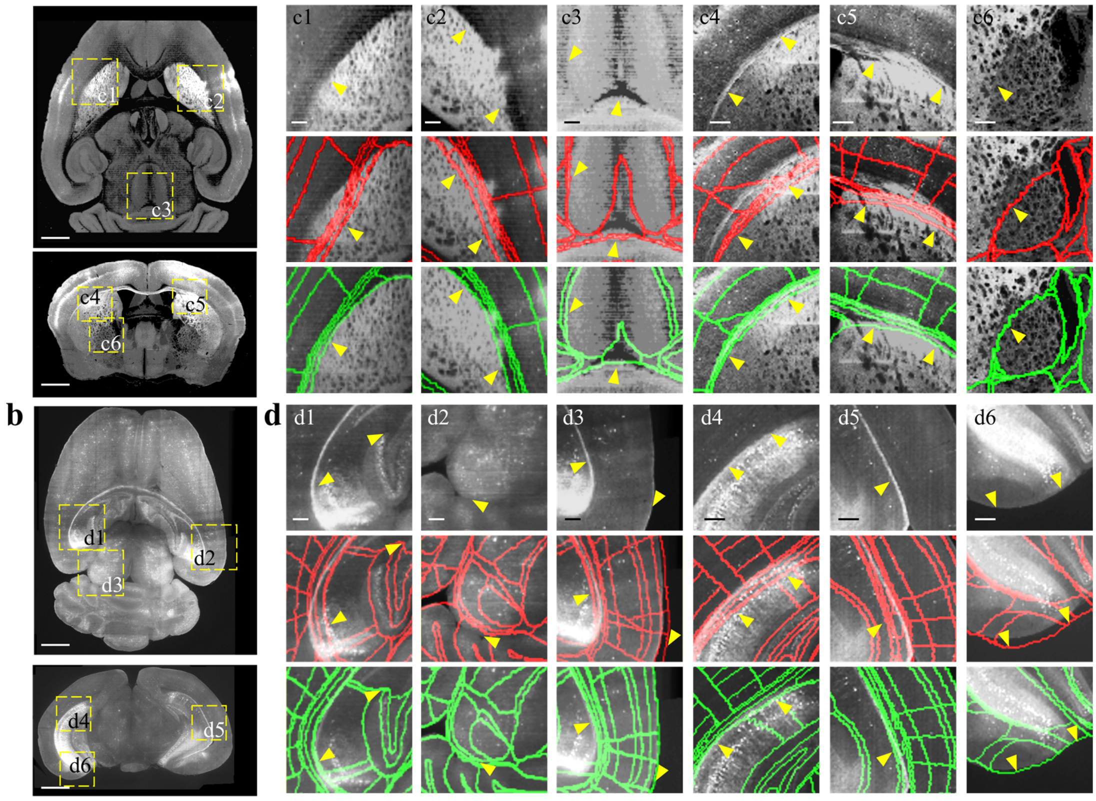


**Supplementary Figure 6: Comparison between BIRDS and conventional single-channel registration. a, b** STP image of intact brain and LSFM image of clarified brain, which are registered by our BIRDS and conventional single-channel method. **c**, **d** Magnified views of six region-of-interests (ROIs, yellow boxes) selected from the horizontal (top in **a, b**) and coronal planes (bottom in **a, b**) in the STP and LSFM brains, respectively. The comparison between BIRDS (green annotation) and single-channel (red annotation) results indicates obviously less inaccuracy by our BIRDS method (yellow arrows). All segmentation results are directly outputted from the programs, without any manual correction. Scale bar, 1 mm for **a, b**, and 250 µm for **c, d.**

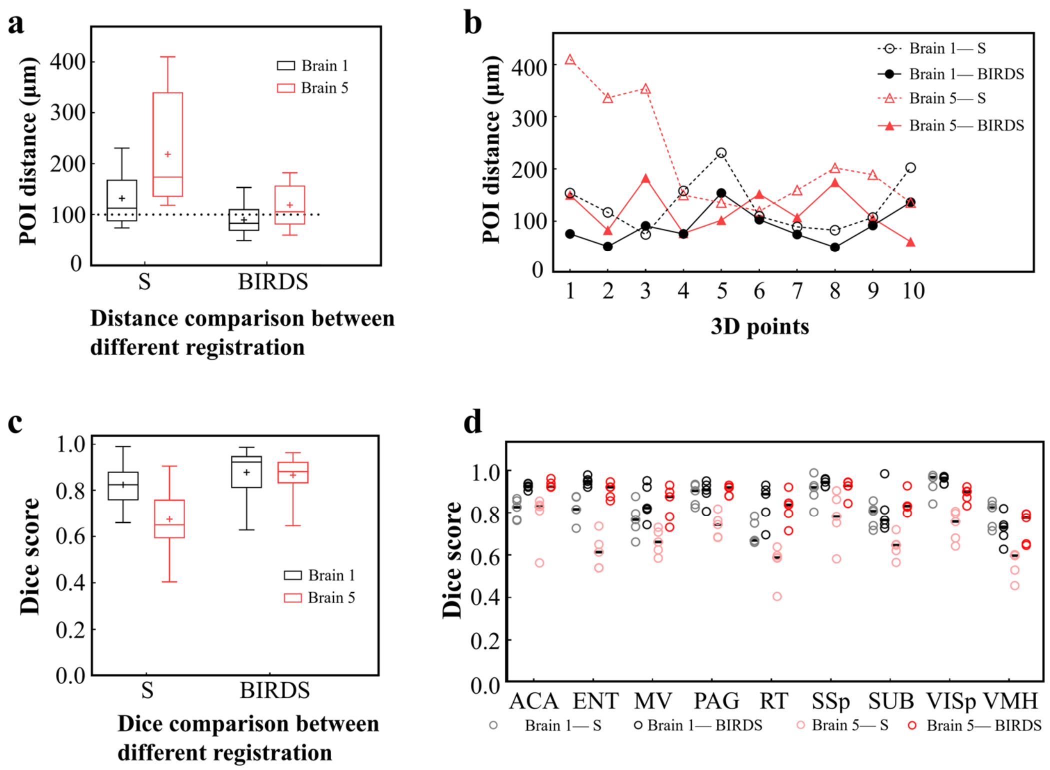


**Supplementary Figure 7: Comparative accuracy analysis of BIRDS and single-channel registration. a** Box diagram showing the distances between the paired POIs (Supplementary Fig. 5) in Allen template and experimental image registered by BIRDS and conventional single-channel registration. Black Brain 1 represents STP image of an intact brain and red Brain 5 represents LSFM image of a clarified brain, which have been also shown in **Supplementary Fig. 6**. The median error distance of 20 pairs of POIs in the two brains registered by BIRDS is ~104 µm, which is compared to ~175 µm error distance by single-channel registration. **b** Line chart specifically showing the distance of each pair of points in two types of datasets by two registration methods. Besides smaller median error distance, here BIRDS registration with including feature information also yields smaller distance variance (solid lines), as compared to single-channel method (dash line). **c** Dice score comparison of two registration methods at nuclei level. 18 regions in two brains (9 regions for each) are selected for the analysis, with results grouped by the datasets. It's clearly shown that the median Dice scores by our method are higher than those by single-channel registration, especially for the highly-deformed clarified brain (Brain 5, 0.65 *vs* 0.881). **d** Dice scores of two registration methods, with results grouped by 9 selected sub-regions (5 planes included for each region), which are ACA, ENT, MV, PAG, RT, SSp, SUB, VISp, and VMH. The black lines indicate the median DICE scores for each regions. Overall, dice scores with averaged median value > 0.9 (calculated by 2 brains) or >0.87 (calculated by 9 regions) were obtained by our BIRDS. These two values are obviously higher than 0.74 and 0.76 by single-channel registration.

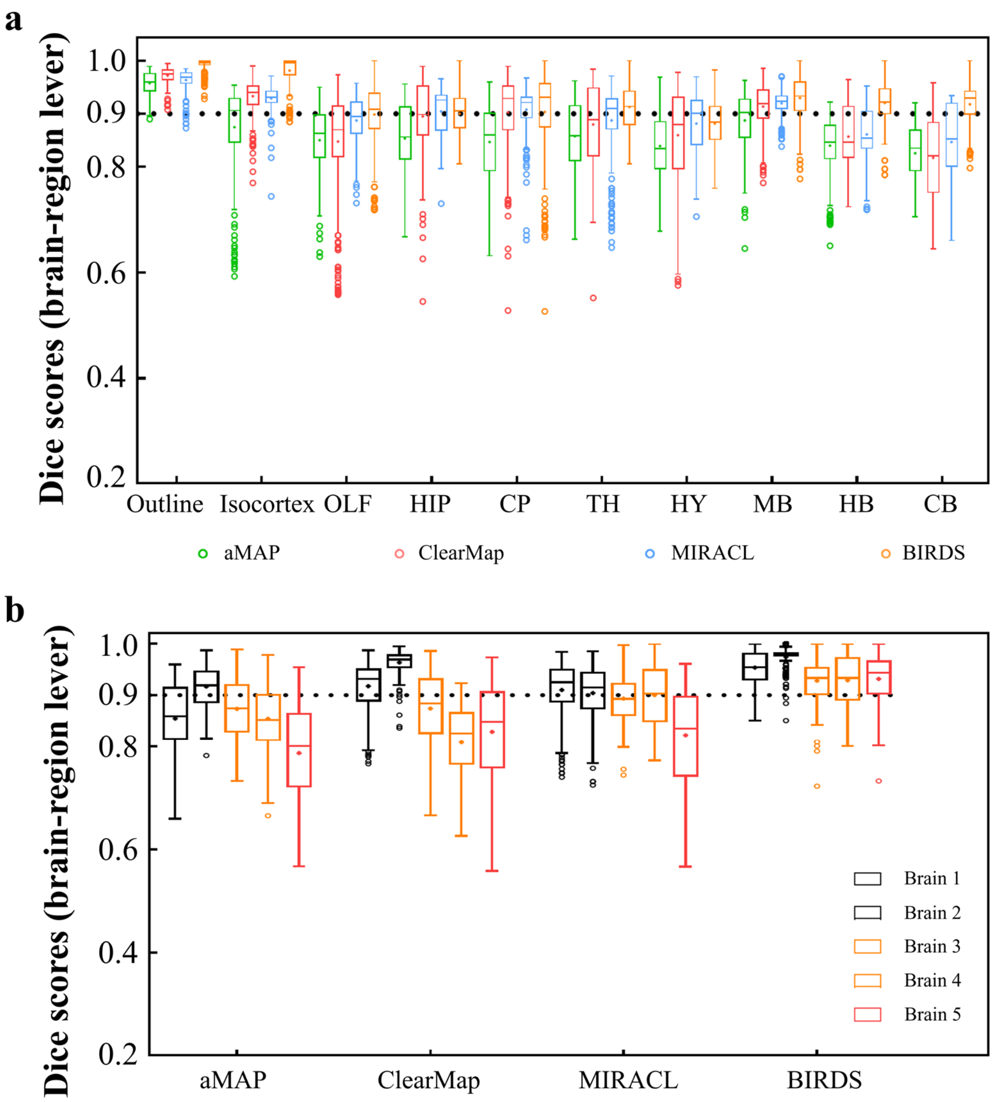


**Supplementary Figure 8: Dice score comparison of nine regions in five brains registered by four registration tools, aMAP, ClearMap, MIRACL and our BIRDS.** The results are grouped by brains in **a**, and regions in **b**. The calculation/comparison is implemented at region level with ~100-µm resolution. When the results are analyzed by brains, BIRDS surpass the other 3 methods most on LSFM dataset #5, with 0.944 median Dice score being compared to 0.801 by aMAP, 0.848 by ClearMap, and 0.835 by MIRACL. At the same time, all the methods perform well on STP dataset #2 with median Dice of 0.919 by aMAP, 0.969 by ClearMap, 0.915 by MIRACL, and 0.977 by our BIRDS. When the results are compared by 9 functional regions, the median values acquired by our BIRDS were also higher than the other three methods. Even the lowest median Dice score by our method is still 0.885 (indicated by black line), which is notably higher than 0.834 by aMAP, 0.82 by ClearMap, and 0.853 by MIRACL.

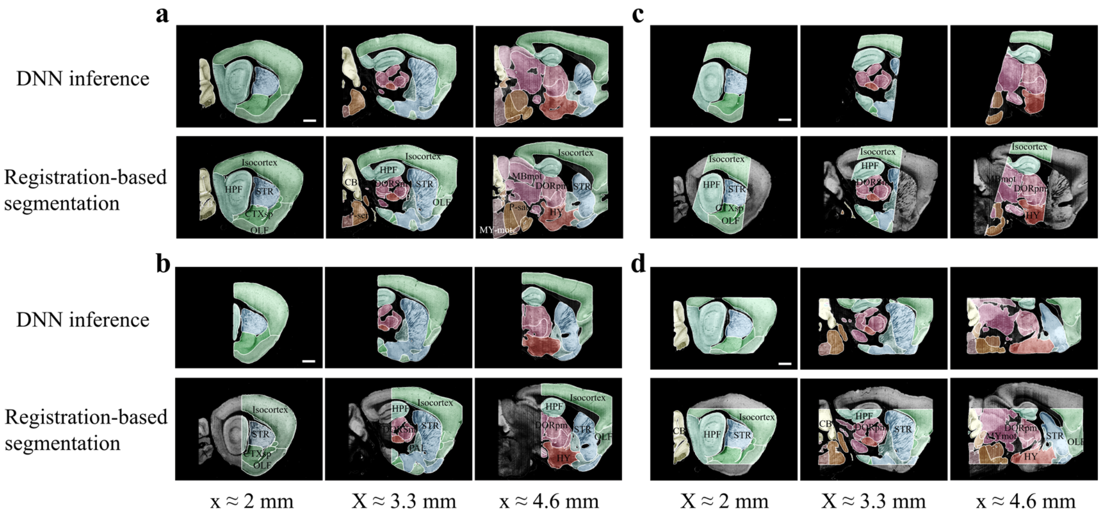


**Supplementary Figure 9: Accuracy comparison between DNN-based and registration-based brain segmentation.** We test the accuracy of DNN-inference-based segmentation through comparison with bi-channel registration results (after manual correction). An entire STP brain and three types of cropped incomplete brain regions, which are the same modes of data selected for network training (**Fig. 2b**), are included for performance validation, with results shown in **a** to **d**. In the comparative analysis of each group of data, we showed the same three sagittal planes (20-μm interval) segmented by DNN inference (upper row) and bi-channel registration (lower row). Eighteen segmented regions compared in the four modes of brain data (Isocortex, HPF, OLF, CTXsp, STR, PAL, CB, DORpm, DORsm, HY, MBsen, MBmot, Mbsta, P-sen, P-mot, P-sat, MY-sen, MY-mot ) have widely verified the sufficient segmentation accuracy by our DNN inference. Scale bar, 1mm.

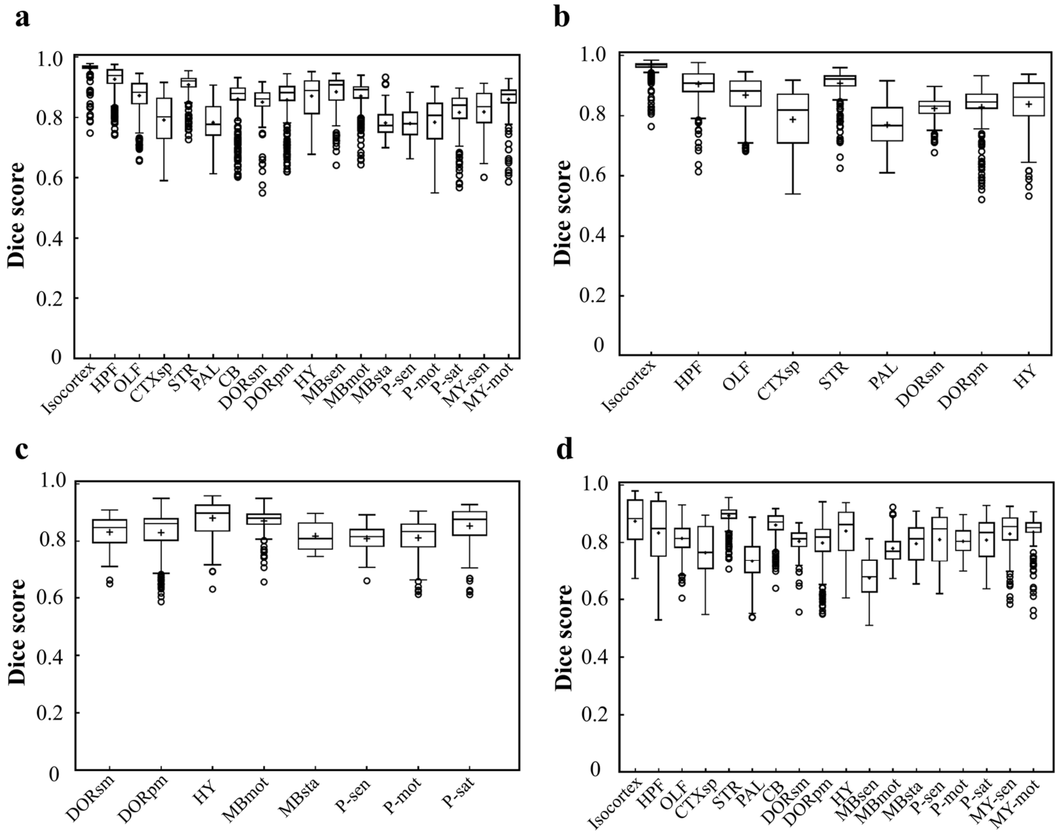


**Supplementary Figure 10: Dice scores of the DNN-segmented regions in abovementioned four types of brains.** Data plots in **a, b, c, d** correspond to segmentation results shown in Supplementary **Fig. 10a, b, c, d,** respectively. The average median value of Dice scores for most of the individual regions in all four brain modes are above 0.8, e.g., Isocortex (~0.941), HPF (~0.884), OLF (~0.854), STR (~0.908), thereby showing a high inference accuracy for most of brain regions. At the same time, the performance of the network for segmenting PAL (~0.761) and MBsta (~0.799) regions remain limited, possibly owing to the large structure variation of these two regions as we move from the lateral to medial images across the sagittal plane.

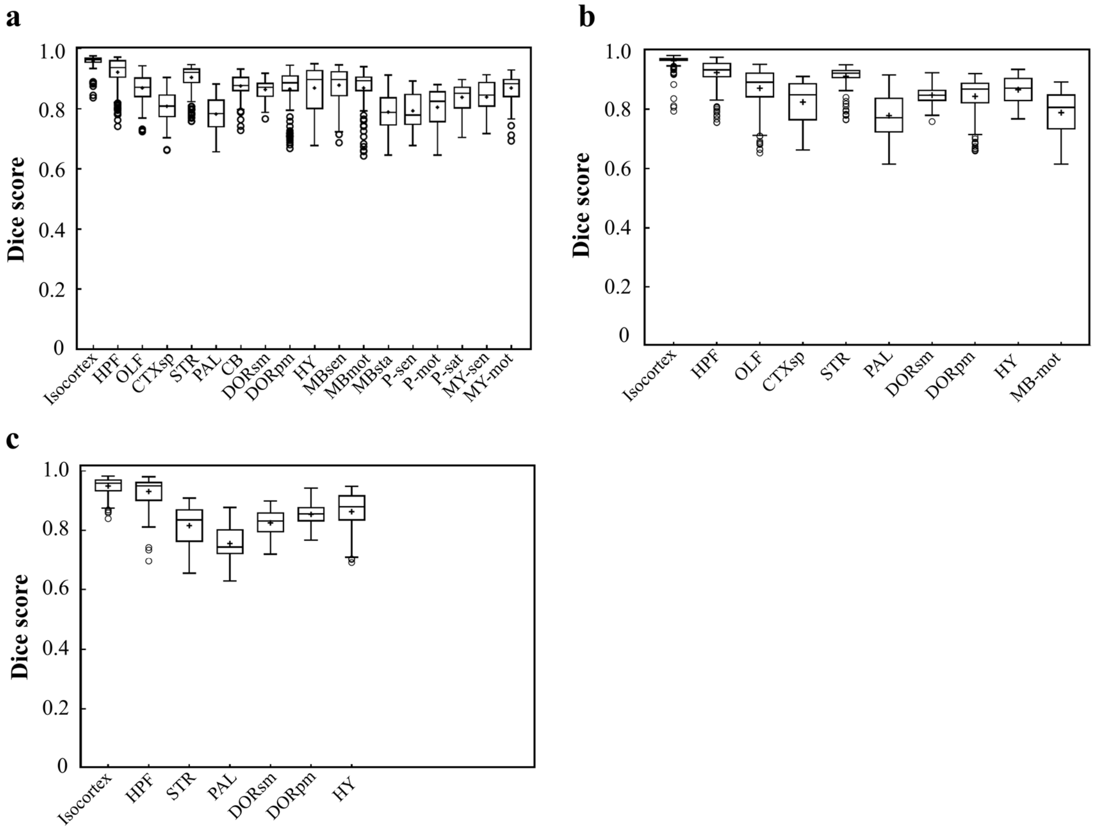


**Supplementary Figure 11: Dice scores of the DNN-segmented regions in three new types of incomplete brains.** These three incomplete brains, also shown in Fig. 4 c, d, e, all look different with the training data modes. Thereby, we can test the real inference capability of the trained DNN for segmenting unfamiliar data. **a** Dice scores of 18 DNN-segmented regions in right hemisphere. The ground-truth references used for calculation are the results obtained by bi-channel registration plus manual correction. The averaged median value of Dice scores was 0.86, with maximum value of 0.965 at Isocortex and minimum value of 0.78 at P-sen. **b** Dice scores of 10 DNN-segmented regions in an irregular cut of half telencephalon. The averaged median value of Dice scores was 0.87, with maximum value of 0.97 at Isocortex and minimum value of 0.771 at PAL. **c** Dice scores of 7 DNN-segmented regions in a small random cut of brain. The averaged median value of Dice scores was 0.86, with maximum value of 0.958 at Isocortex and minimum value of 0.745 at PAL.

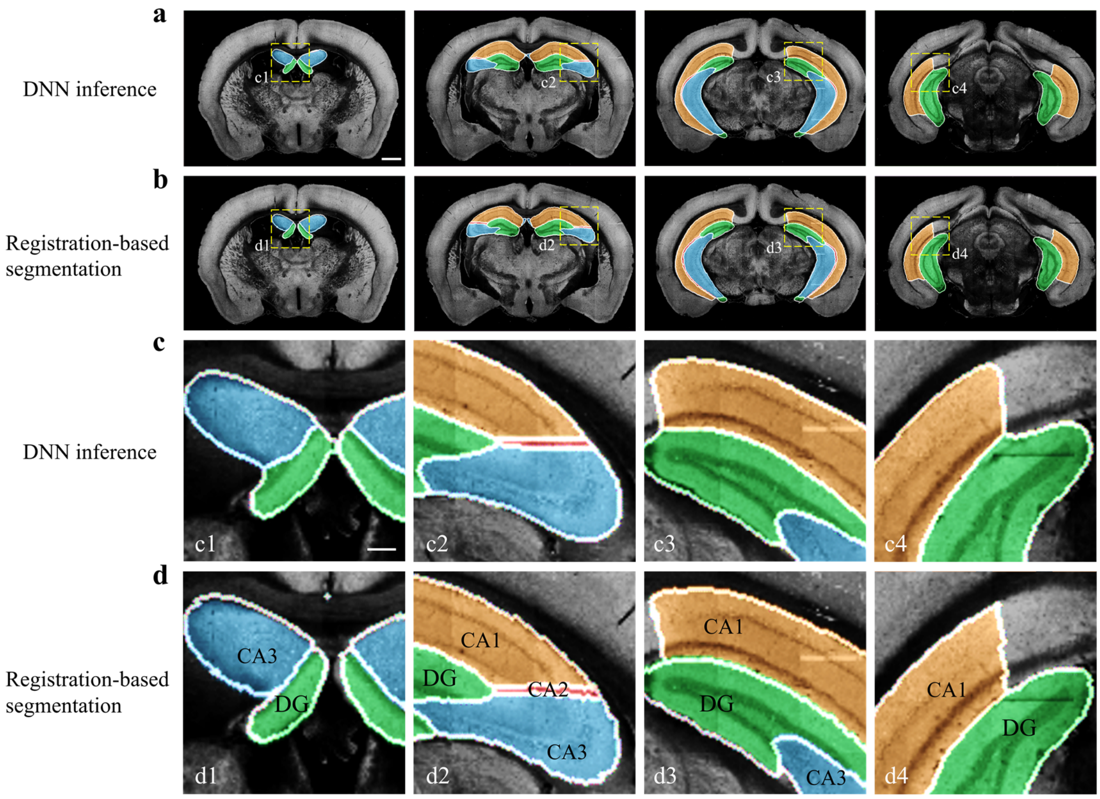


**Supplementary Figure 12: Performance of DNN inference for segmenting fine sub-regions specifically in the hippocampus.** Four coronal planes containing hippocampal sub-regions of CA1, CA2, CA3, and the DG are shown and compared to the bi-channel registration results. **a, b,** Color-rendered segmentations of CA1, CA2, CA3, and DG by network inference and bi-channel registration (with manual correction), respectively. Scale bar, 1mm. **c, d,** Magnified views of the selected small regions (boxes) revealing the highly similar segmentation results by DNN inference and bi-channel registration. Scale bar, 250 µm. The averaged median value of Dice scores was 0.878, with maximum value of 0.96 at CA1 and minimum value of 0.70 at CA2.

**a**

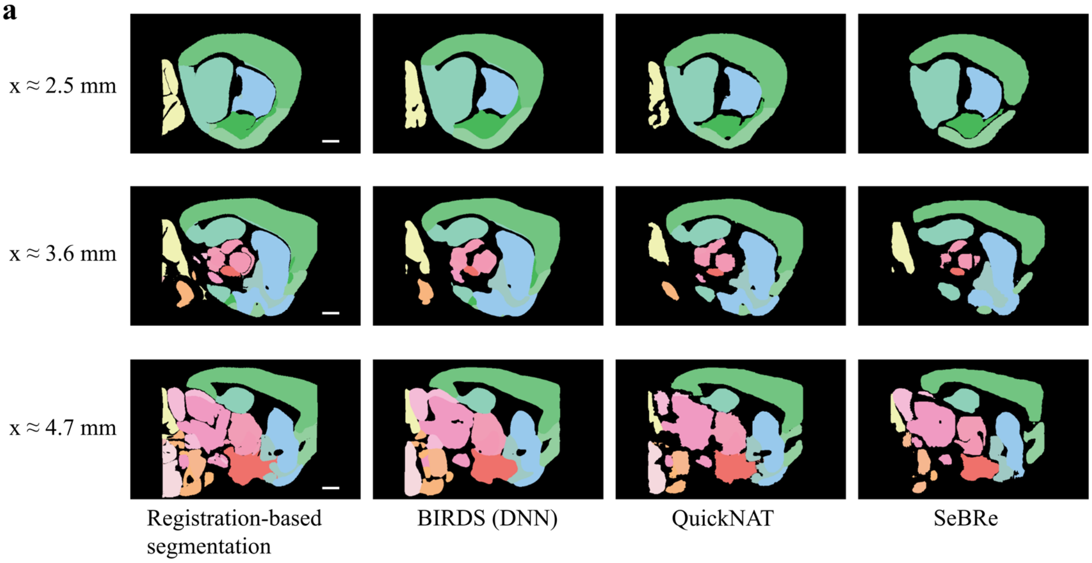


**b**

| | BIRDS (DNN) | QuickNAT | SeBRe | | BIRDS (DNN) | QuickNAT | SeBRe |
|---|---|---|---|---|---|---|---|
| Isocortex | 0.9647 * | 0.9178 | 0.858 | HY | 0.8896 * | 0.8289 | 0.8148 |
| HPF | 0.9381 * | 0.909 | 0.8755 | MBsen | 0.9092 * | 0.8378 | 0.8005 |
| OLF | 0.8705 * | 0.858 | 0.8558 | MBmot | 0.892 * | 0.8573 | 0.8196 |
| CTXsp | 0.8017 | 0.8411 * | 0.8143 | MBsta | 0.774 | 0.8148 * | 0.7972 |
| STR | 0.9128 * | 0.8767 | 0.8632 | P-sen | 0.7799 * | 0.7535 | 0.7383 |
| PAL | 0.7771 | 0.7823 * | 0.7379 | P-mot | 0.8069 * | 0.7939 | 0.7747 |
| CB | 0.8787 * | 0.8152 | 0.7169 | P-sat | 0.841 * | 0.8118 | 0.7171 |
| DORsm | 0.8623 * | 0.8362 | 0.8148 | MY-sen | 0.8365 * | 0.8144 | 0.8012 |
| DORpm | 0.8821 * | 0.8311 | 0.7801 | MY-mot | 0.8768 * | 0.8289 | 0.681 |

* The best performance on the given brain region

**Supplementary Figure 13: Comparison of the performance of DNN inference by three deep learning-based brain region segmentation techniques, QuickNAT, SeBRe and our BIRDS (DNN). a**. BIRDS (Bi-channel registration) and DNNs-inference-based segmentation results of an entire STP brain. Scale bar, 1mm. In the comparative analysis of each method, we showed the same three sagittal planes (20-µm interval) segmented by DNN inference (BIRDS, QuickNAT, SeBRe) and bi-channel registration. Eighteen regions (Isocortex, HPF, OLF, CTXsp, STR, PAL, CB, DORpm, DORsm, HY, MBsen, MBmot, Mbsta, P-sen, P-mot, P-sat, MY-sen, MY-mot ) segmented by the three deep learning-based methods have widely verified the sufficient segmentation accuracy by DNN inference. In most of brain regions the performance of BIRDS (DNN) is the best among the three (*), except CTXsp, PAL, and MBsta (QuickNAT is better than BIRDS (DNN) and SeBRe).

#### Supplementary Table 1: Data size and computational cost at different BIRD stages

| Stage | Dataset | Resolution ($\mu m^3$) | Volume ($pixel^3$) | Image format | Data size (GB) | Transform time (min) |
|---|---|---|---|---|---|---|
| 1: image preprocessing (segmented downsampling) | raw image （Brain 1） | 1*1*10 | 11972*7536*1075 | 16bit | 180 | 5~10 |
| | Allen Reference Atlas | 20*20*100 | 570*400*132 | RGB | 0.336 | |
| 2: Registration | fix image (the result of image preprocessing) | 20*20*20 | 570*400*660 | 8bit | 0.137 | 15~20 |
| | move image (average template) | | | 8bit | 0.137 | |
| | registration result | | | 8bit | 0.137 | |
| 3: Transformation (the parameter from registration) | annotation file | 20*20*20 | 570*400*660 | 32bit | 0.561 | 0.5 |
| | annotation file（after registration） | | | 32bit | 0.561 | |
| 4： 3D digital framework | 3D digital map | 1*1*10 | 11972*7536*1075 | 16bit | 360 | 20~30 |
